# CDK7 is a Novel Therapeutic Vulnerability in Fibrolamellar Carcinoma

**DOI:** 10.1101/2023.04.22.537934

**Authors:** Manabu Nukaya, Crystal Cafferty, Katerina Zahed, Isabelle Yun, David P. Al-Adra, Noor A. Kazim, Alaa R. Farghli, Marina Chan, Jeremy D. Kratz, Mark E. Berres, Andrew Yen, Taranjit S. Gujral, Praveen Sethupathy, Christopher A. Bradfield, Sean M. Ronnekleiv-Kelly

## Abstract

Fibrolamellar carcinoma (FLC) is a rare and lethal cancer that afflicts young individuals. The tumor arises in the background of a healthy liver, and patients typically present with advanced cancer at the time of diagnosis. Unfortunately, for these patients with advanced or recurrent cancer, no proven systemic therapies exist resulting in only 30-45% of patients surviving to 5 years. Investigations into the molecular underpinning of FLC have revealed a unique gene fusion between heat shock protein 40 (*DNAJB1*) and the catalytic subunit alpha of protein kinase A (*PRKACA*), leading to the formation of an oncoprotein (DNAJ-PKAc) that retains kinase activity and is a proven tumor-causing event in FLC. To uncover potential therapeutic targets, we engineered an FLC cell line by introducing the *DNAJB1-PRKACA* oncogene rearrangement into human hepatocellular cells using CRISPR/Cas9. We identified aberrant cell cycle progression, and follow-up molecular analysis revealed evidence of enhanced cyclin dependent kinase 7 (CDK7) activation in the *DNAJB1-PRKACA* expressing FLC cells. These findings were confirmed in human samples of FLC. In turn, targeting CDK7 with selective inhibitors demonstrated efficacy in several patient-derived models of FLC, with minimal toxicity to normal liver. Collectively, this work uncovers a novel candidate therapeutic vulnerability in FLC.

## INTRODUCTION

Fibrolamellar carcinoma (FLC) is a rare and lethal cancer that afflicts young individuals, and accounts for approximately 1% of liver cancers.^1-4^ The tumor arises in the background of a normal liver, and patients typically present with advanced cancer at the time of diagnosis.^2, 5^ This is consequent to the absence of known risk factors or tumor markers that can precipitate screening or prevention.^1, 6 7-9^ While surgery offers the best chance for long-term survival, most patients experience cancer recurrence after surgical resection.^1, 5^ Unfortunately, for these patients with advanced or recurrent cancer, no proven systemic therapies exist, and only 30-45% of patients survive to 5 years.^1, 6^ When considering that the peak age of diagnosis is 22 years,^2^ the substantial impact on life-years lost underscores the devastating nature of this cancer. Thus, there remains a critical need to improve outcomes for patients with FLC through systemic treatment.

Investigations into the molecular underpinning of FLC have revealed a unique gene fusion between heat shock protein 40 (*DNAJB1*) and the catalytic subunit alpha of protein kinase A (*PRKACA*).^6^ The consequent gene fusion (*DNAJB1-PRKACA*) is the proven driver mutation of FLC in human (vast majority) and mouse, and the resultant chimeric protein (DNAJ-PKAc) is only present in tumor cells (somatic event).^5, 10, 11^ The DNAJ-PKAc oncoprotein is therefore a viable target in FLC. This was demonstrated in recent work from the group that discovered the oncogene fusion in FLC. They uncovered that not only is *DNAJB1-PRKACA* sufficient for FLC development, but that it is required for tumor cells to survive.^12^ They subsequently showed the efficacy in targeting *DNAJB1-PRKACA* with shRNA in human-relevant models, but translating this approach will require minimizing off-target effects of native PKA and ensuring effectiveness *in situ*.^12^ Another group exploited the FLC-specific nature (*i.e*., exclusively in tumor cells) of the immunogenic neoepitopes from the DNAJ-PKAc chimera to pioneer an innovative immune targeting strategy.^13^ However, limitations exist with a vaccine / immunotherapy-based strategy (*e.g*., patients with autoimmune disorder, immunotherapy-induced toxicity, or lack of induced immune response).^14-16^ Thus, while highly promising strategies to target the initiating and sustaining event in FLC (*DNAJB1-PRKACA*) are being developed, certain barriers remain.

Meanwhile, another landmark study showed that drug-repurposing (*i.e*., not FLC specific) led to a subset of effective tumoricidal treatments (*e.g.*, Irinotecan) in patient-tumor-derived xenografts of FLC.^17^ Despite the effectiveness demonstrated by a subset of drug treatments, there is still uncertainty surrounding the reason why certain drug compounds were effective in the preclinical models of FLC; the connection between DNAJ-PKAc fusion and drug treatment effects remains unclear. Concordantly, there have been concerted efforts to understand how *DNAJB1*-*PRKACA* drives tumor development and mechanisms of treatment resistance so that new and effective therapies can be identifed.^11, 18^ For instance, the chimeric protein demonstrates enhanced PKA activity compared to native PKA in response to cAMP,^6^ but simply inhibiting PKA is not a viable therapeutic option in FLC given the critical nature of PKA in normally functioning cells (*i.e.,* no therapeutic window).^12^ Additionally, PKA overactivity is unlikely to be the sole cause of its carcinogenic effects.^18^ In support of this assertion, the DNAJ-PKAc fusion oncoprotein was found to disrupt type I regulatory subunit (RIα) compartmentalization of cAMP, causing dysregulated cAMP-dependent signaling.^19^ This results in activation of proteins not typically phosphorylated by native PKA.^18^ While the precise signaling cascade remains unknown, recent seminal work has illuminated a transcriptional molecular endpoint. Specifically, the fusion oncoprotein causes aberrant remodeling of the epigenetic landscape and emergence of FLC specific super enhancers to ultimately drive expression of FLC-specific targets like SLC16A14, LINC00473, miR-375 and miR-10b.^11, 20, 21^ Thus, these too represent highly promising biological targets in FLC.

Yet, a critical gap exists in understanding how the tumor-driver mutation (*i.e*., *DNAJB1-PRKACA*) transmits signals to ultimately promote super enhancer driven gene expression in FLC. Determining this pathogenesis has proven challenging due to the limited models available for mechanistic studies, and the difficulty in parsing out PKA-specific effects versus DNAJ-PKAc driven pathology. Addressing this gap could uncover novel targets in FLC and reveal unique therapeutic vulnerabilities, which could be coupled with emerging strategies to enhance tumor-killing effect.

Therefore, we created an FLC cell line by introducing the *DNAJB1-PRKACA* oncogene rearrangement into a human hepatoblastoma cell line (HepG2). The purpose of this cell line was to use it as a tool to elucidate the DNAJ-PKAc driven pathogenic mechanism; any pertinent findings were then confirmed in patient-derived models or human tissue. We identified aberrant cell cycle progression due to the fusion oncogene and found gene expression signatures concordant with the transcriptional dysregulation seen in human FLC. RNA sequencing analysis also revealed evidence of enhanced cyclin-dependent kinase 7 (CDK7) activation, and this target proved to have concurrent involvement in cell-cycle regulation as well as super enhancer driven gene expression. In turn, we found that CDK7 is a novel regulator of FLC specific super enhancer driven genes, and a therapeutic vulnerability in human FLC.

## RESULTS

### Engineering DNAJB1-PRKACA expressing cells to evaluate FLC pathogenesis

We generated the *DNAJB1-PRKACA* gene fusion in a hepatoblastoma cell line derived from a 15-year-old male (HepG2),^22^ using dual-guide RNA/Cas9 that targets the intronic region between exon 1 and 2 of *DNAJB1* and the intronic region between exon 1 and 2 of *PRKACA* (**Figure 1A-B**). Genomic DNA PCR and sequencing across the breakpoint established presence of the *DNAJB1-PRKACA* gene fusion in six separate clones (**Figure 1C-D**). We confirmed mRNA expression of *DNAJB1-PRKACA* and performed sequencing of the mRNA transcripts to ensure development of the expected fusion transcript (**Figure 1D, 1E**). Protein analysis demonstrated presence of DNAJ-PKAc and native PKA, which is the expected pattern of protein detection in human FLC (**Figure 1F**).^6^ Based on expression levels of native genes (e.g., *PRKACA*) and FLC-specific markers (e.g., *SLC16A14* and *LINC00473*), we selected clone H33 (HepG2^DNAJB1-PRKACA^) for subsequent assays (**Figure 1E**). Another clone, H12 (HepG2^DNAJB1-PRKACA^), also met the same rigorous validation metrics and was utilized for confirmatory tests.

**Figure 1.**
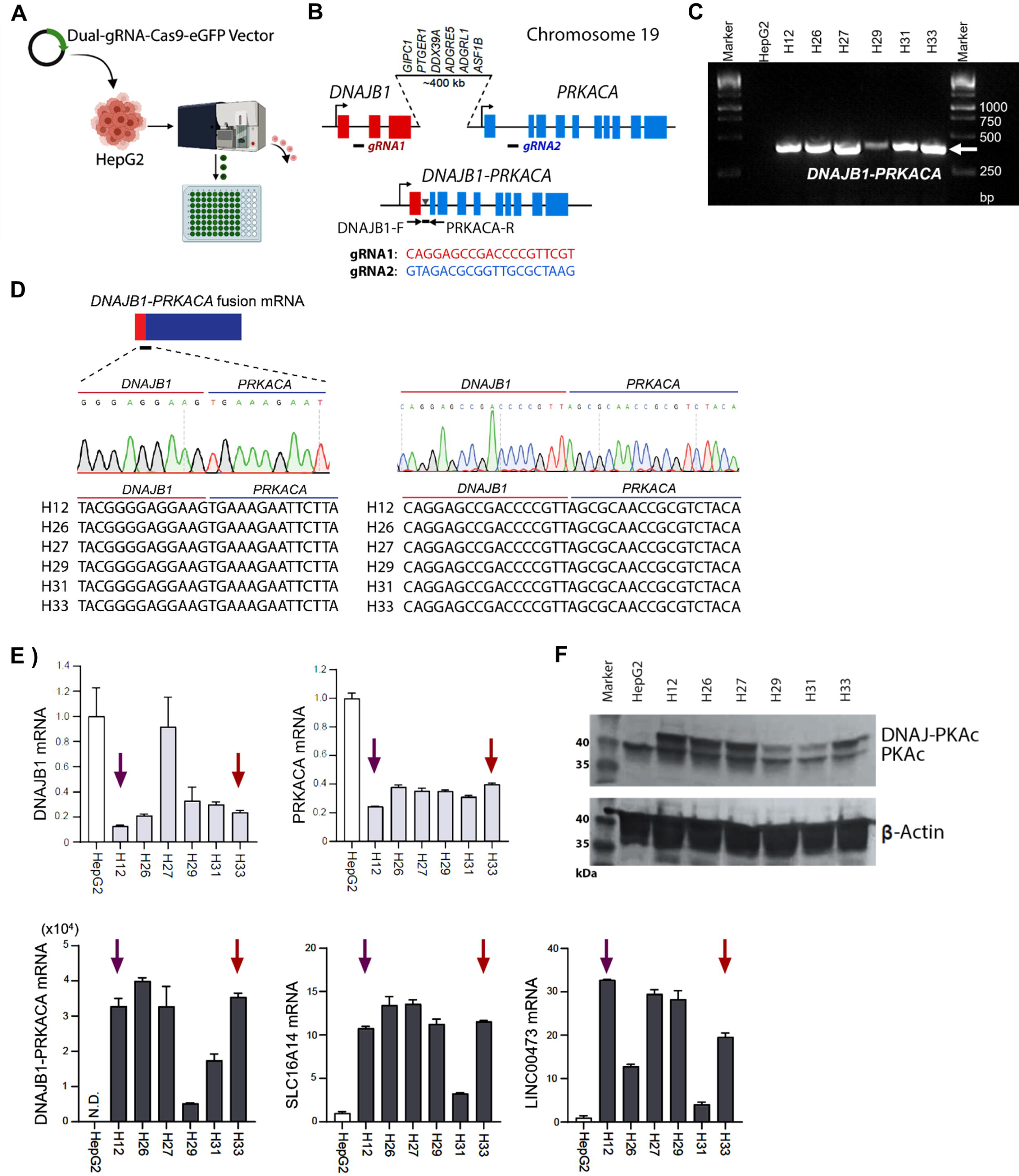
*Generation of a model to understand pathogenic mechanisms of DNAJB1-PRKACA in FLC*. Parent hepatoblastoma cells (HepG2) were transfected with a dual-guide RNA-CAS9 plasmid containing an eGFP tag, to allow for cell sorting of HepG2 cells that took up the plasmid. Each cell was deposited in a single well of a 96-well plate and clonally expanded (**A**). The guide RNAs (gRNA1: CAGGAGCCGACCCCGTTCGT, gRNA2: GTAGACGCGGTTGCGCTAAG) directed CAS9 to induce a double-strand break at intron 1 of *DNAJB1* and intron 1 of *PRKACA*, resulting in 400kb deletion and chromosomal rearrangement to generate the *DNAJB1-PRKACA* oncogene fusion (**B**). After expansion, genomic DNA from each clone was assessed via PCR for presence of *DNAJB1-PRKACA* gene fusion, and six clones manifested appropriately sized PCR products (DNA bands) pertaining to the forward and reverse primers across the breakpoint (**B-C**). The PCR product from each clone and *DNAJB1-PRKACA* mRNA transcript (across the breakpoint) was sequenced to determine the precise sequence by CRISPR gene editing (**D**). Assessment of expression levels of native genes *DNAJB1* and *PRKACA* (light bars), as well as FLC-specific genes including *DNAJB1-PRKACA* fusion, *SLC16A14* and *LINC00473* (dark bars) was performed. Shown is relative fold-change (error bars represent standard deviation) compared to parent HepG2 cells, with four biological replicates per clone (**E**). Clone samples and parent HepG2 cells were also evaluated for generation of fusion oncoprotein (DNAJ-PKAc) and native PKA (β-Actin control) (**F**). Clone selection for subsequent assays was based on these rigorous validation metrics (**A-F**) yielding H33 (red arrow) and H12 (purple arrow) as the selected clones.

### DNAJB1-PRKACA drives accelerated cell cycle progression

H33 cells (possessing the *DNAJB1-PRKACA* gene fusion) demonstrated similar proliferation rates as parent HepG2 cells, which is concordant with prior work evaluating the impact of introducing *DNAJB1-PRKACA* into a liver cancer cell line (**Figure 2A**).^12^ Yet, when performing a cell cycle analysis, we found that H33 cells showed a significant increase in the percent of cells in G2/M (40.4% vs 26.4%, *p* = 0.003) and significant decrease in percent of cells in G0/G1 (32.5% vs 47.6%, *p* = 0.0004) compared to parent HepG2 cells (**Figure 2B**). This was confirmed in a time-course experiment in which H33 and HepG2 cells were synchronized by serum-free conditions and subsequently evaluated for cell cycle progression after return to serum-containing culture media. H33 cells demonstrated an earlier drop in percent of cells in G0/G1, corresponding to an earlier peak in S-phase (12hr, *p* < 0.0001) and a more substantive progression to G2/M (24hr, *p* < 0.0001) than HepG2 cells (**Figure 2C**). Given the higher replicative capability, we consequently assessed levels of apoptosis. After plating equivalent number of cells, H33 cells demonstrated a significant increase in apoptosis (*p* < 0.001) over a four-day period as measured by caspase 3/7 and caspase 8 activity (**Figure 2D**). To confirm these findings, we tested a separate clone (H12) that possessed the *DNAJB1-PRKACA* fusion and the H12 cells demonstrated nearly identical increases in caspase 3/7 and caspase 8 activity (*p* < 0.001). To validate the results, poly (ADP-ribose) polymerase (PARP1) and cleaved-PARP1 protein were evaluated in HepG2 and H33 cells using two separate antibodies that i) detect full length PARP1 and cleaved PARP1, and ii) detect only cleaved PARP1 protein. PARP1 is a primary protein target of caspase-3, and cleavage of PARP1 is a marker of cellular apoptosis.^23, 24^ Cleaved PARP1 was increased in H33 cells compared to HepG2 cells, corroborating the heightened caspase 3/7 and caspase 8 activity (**Figure 2D**). Collectively, this demonstrates that there is an increase in cell cycle progression in *DNAJB1-PRKACA* expressing H33 cells versus parent HepG2 cells, with an associated rise in apoptosis that balances proliferation rates.

**Figure 2.**
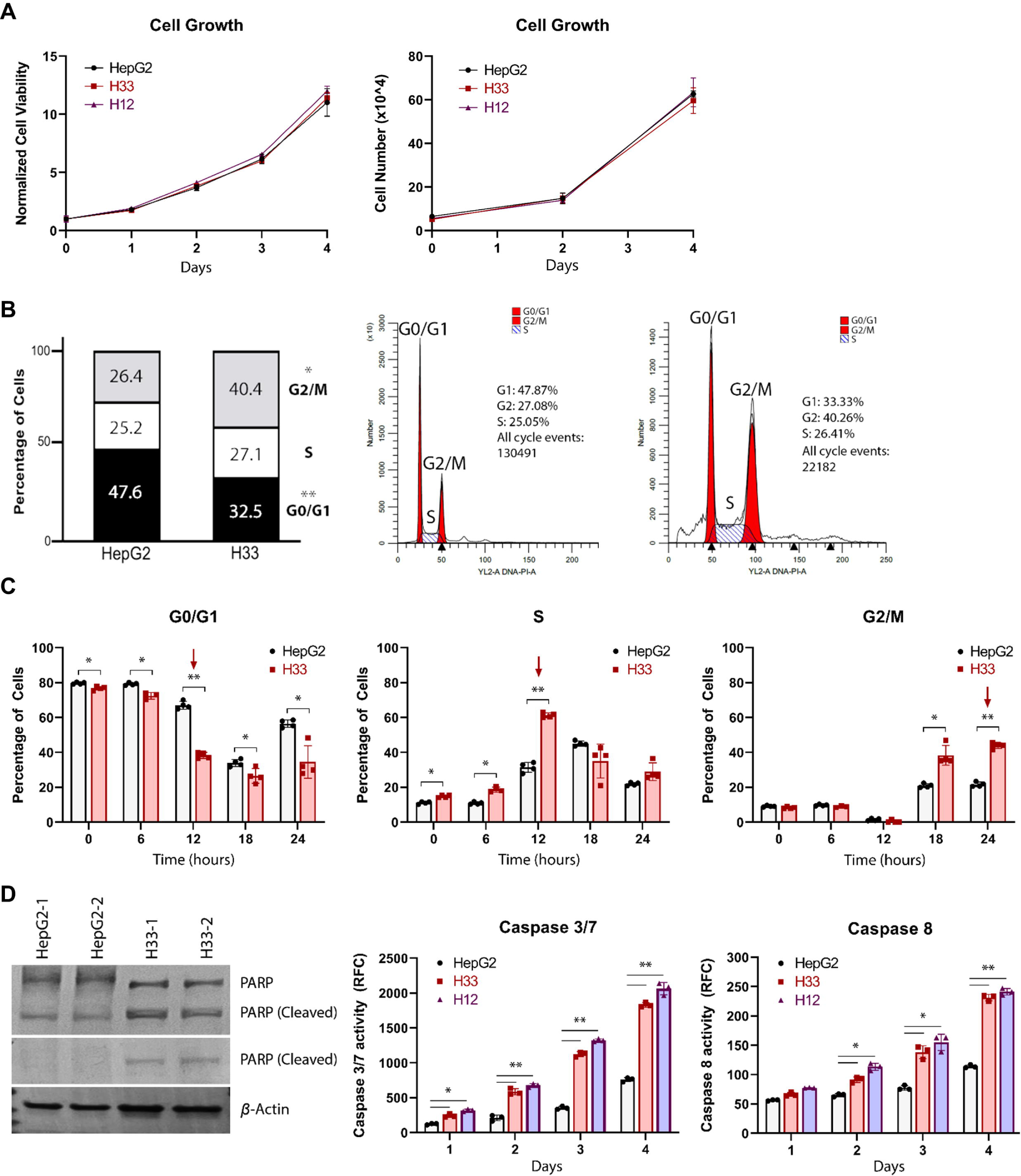
*DNAJB1-PRKACA induces changes in cell cycle progression and apoptosis*. Cell proliferation of H33 cells and H12 cells (both possessing the *DNAJB1-PRKACA* gene fusion) were measured and compared to HepG2 cells over 4 days, with viability assessed by normalized luminescence (normalized to luminescence value at day 0 after plating identical number of cells). As a separate confirmatory test, HepG2, H33 and H12 cells were plated and counted via hemocytometer. After two days and four days, the cell number was quantified via hemocytometer and reported as total cell number (x10^4^) (**A**). Using propidium iodide (PI) staining and flow cytometry, the percent of cells in G0/G1 (\*\**p* = 0.0004), S-phase and G2/M (\**p* = 0.003) were evaluated and compared between HepG2 and H33 cell lines. Shown also are representative flow cytometry graphs for HepG2 (left) and H33 (right) depicting the percent of cells in each phase, with the first peak pertaining to G0/G1, the second peak pertaining to G2/M, and the region in between representing S-phase (**B**). HepG2 and H33 cells were cultured in serum-free media to synchronize the percent of cells in G0/G1 versus those undergoing active cell division. Serum containing media was then added (Time 0) and using flow cytometry, the percent of cells in G0/G1, S-phase, and G2/M were calculated every 6 hours for 24 hours. The red arrow demarcates the timing in each phase where the greatest difference occurred (\**p* < 0.05, \*\**p* < 0.0001) (**C**). Assessment of apoptosis in HepG2 and H33 was measured by caspase 3/7 activity and caspase 8 activity over a period of 4 days, with an additional *DNAJB1-PRKACA* expressing clone (H12) evaluated to confirm the findings. Significance indicates H33 and H12 compared to HepG2 (\**p* < 0.01, ** *p* < 0.001). To validate the results, PARP and cleaved-PARP (main cleavage target of caspase-3) protein were evaluated in HepG2 and H33 cells. The first antibody detects the endogenous (116kDa) PARP1 protein and the carboxy-terminal domain of cleaved-PARP1 (89kDa), while the second antibody only detects the carboxy-terminal cleaved-PARP1 fragment (89kDa) and not full length PARP1 or other isoforms (**D**). Each of the experiments (B-C) were performed with four biological replicates per line, while (A) included six biological replicates and (D) included three biological replicates per line.

### Activated CDK7 in cells expressing DNAJB1-PRKACA and human FLC samples

To gain insight into how *DNAJB1-PRKACA* causes dysregulation of the cell cycle we performed RNA sequencing of H33 (n = 6) and HepG2 (n = 6) cell samples and differential expression analysis. We assessed *SLC16A14* to ensure concordance with human FLC and with our RT-PCR data (from clone selection) and found mean expression levels of 1.83 in the HepG2 samples versus 552 in the *DNAJB1-PRKACA* expressing H33 samples (FDR *q* = 0). This was confirmed in the H12 clone (n = 6) (**Figure 3A**).^11^ *LINC00473* was not detected by RNA sequencing (likely due to an annotation issue), so we performed a repeat qPCR to confirm significant upregulation (181-fold H33 vs HepG2, *p* = 0.0007) (**Supplemental Figure 1**). Further, we performed KEGG pathway analysis of differentially expressed genes between the H33 and HepG2 cells, and compared to the enriched pathways from differentially expressed genes between human FLC versus normal liver (from our team’s prior RNA sequencing dataset).^25^ We found strong overlap of enriched pathways including six of the thirteen enriched pathways identified in the human samples (**Supplemental Figure 2**). Interestingly, two of our top differentially expressed genes were secretogranin II (*SCG2*) and serpin family C member 1 (*SERPINC1*), which were vastly upregulated (log2 fold-change = 14.1, FDR *q* <0.0001) and downregulated (log2 fold-change = -12.5, FDR *q* < 0.0001) in H33 vs HepG2, respectively (**Figure 3A**). When evaluating published tumor-normal data from two separate prior landmark studies,^11, 17^ *SCG2* is substantially upregulated and *SERPINC1* – coding for antithrombin (AT) protein – is markedly downregulated in FLC compared to normal liver across all datasets. Using our team’s prior RNA sequencing dataset of FLC tumor (n=35) and normal (n=10), we found substantial upregulation of *SCG2* (mean expression [standard deviation]: 573.4 [1061] vs 8.0 [14.8], *q* <0.0001) and downregulation of *SERPINC1* (9327 [10853] vs 142675 [52639], *q* <0.0001) in tumor (FLC) versus normal tissue (**Figure 3A**).^25^ The role of SCG2 and AT in FLC is currently unclear, but given the consistent identification of substantive differential expression of their respective genes, these may warrant further exploration.

**Figure 3.**
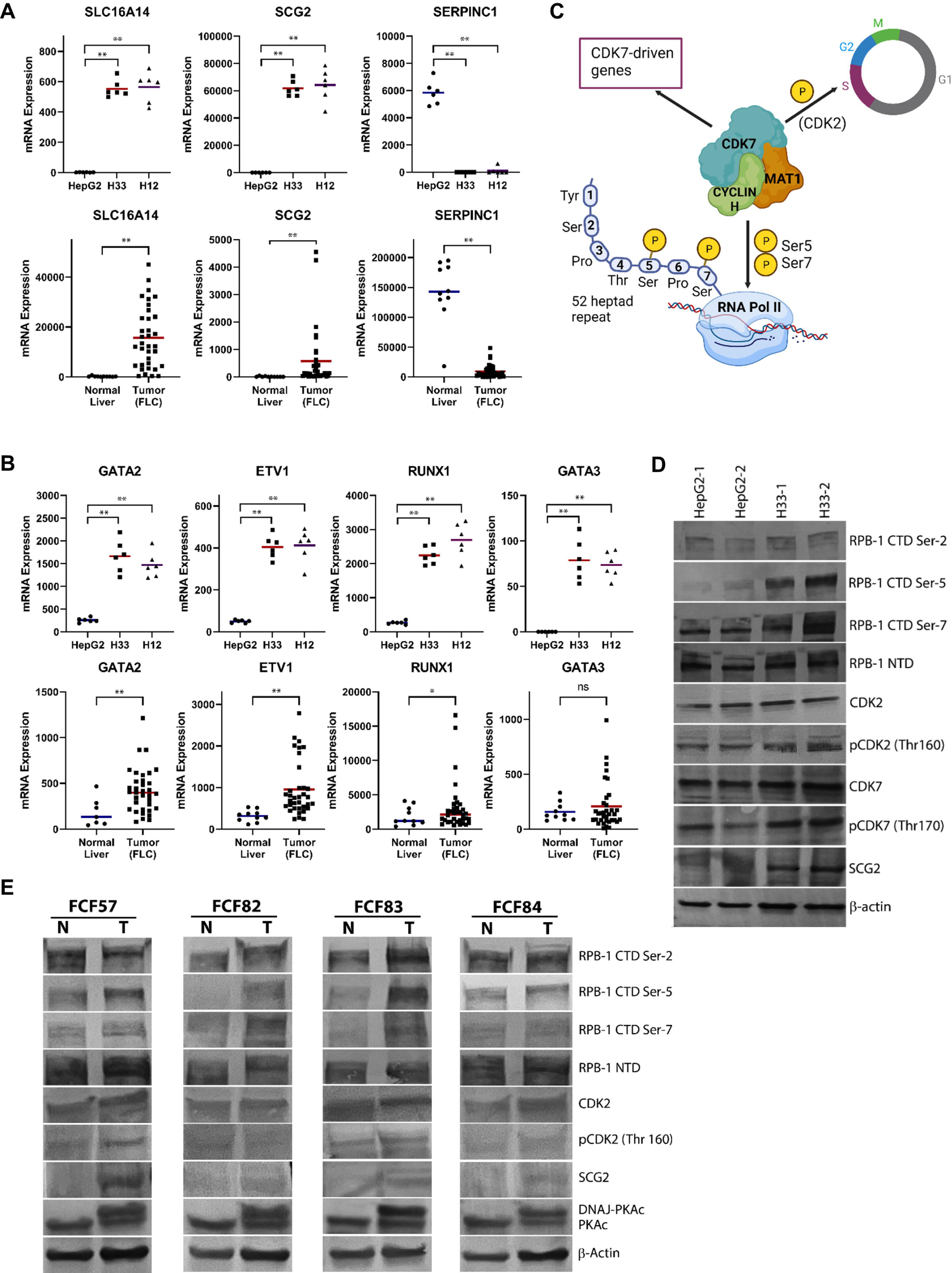
*RNA sequencing and protein analysis reveals CDK7 pathway activation*. RNA sequencing of HepG2 cells and H33 cells (six biological replicates per group) were evaluated for transcriptional changes that have been identified consistently across human tumor-normal FLC datasets, including *SLC16A14*, *SCG2* and *SERPINC1*. Findings from the H33 clone were confirmed in a separate *DNAJB1-PRKACA* expressing clone (H12). mRNA expression of the same targets in human tumor (n = 35) and normal liver (n = 10) samples were assessed (\*\**q* < 0.0001) (**A**). CDK7 regulated genes were evaluated for differential expression in HepG2 vs H33 cells, and findings were validated with the H12 clone (*FDR *q* < 0.0001). The same CDK7 regulated genes were evaluated in human tumor (n = 35) vs normal liver (n = 10) samples (\**q* = 0.018, \*\**q* < 0.001) (**B**). In addition to upregulation of known CDK7 regulated genes, CDK7 activation results in phosphorylation of prime targets including CDK2 (cell cycle) and RNA Polymerase II serine-5 and serine-7 (transcription) [Image created with Biorender.com] (**C**). To confirm CDK7 activation in *DNAJB1-PRKACA* expressing H33 cells vs parent HepG2 cells, protein analysis for phosphorylated RNA polymerase II (serine-2, serine-5, serine-7) was performed. RNA polymerase II (RPB-1) n-terminal domain (NTD) serves as a total RPB-1 control. Additionally, analysis of another prominent CDK7 target – Threonine 160 (Thr160) on CDK2 [pCDK2 (Thr160)] – was assessed. Shown are two separate biological replicates for each line (**D**). To substantiate evidence for CDK7 activation in human samples of FLC, individual patient tumors with matched normal liver (FCF 57, 82, 83 and 84) were evaluated for phosphorylated RPB-1 (Ser-2, -5, -7) and phosphorylated CDK2 (Thr160 pCDK2). Secretogranin II (SCG2) protein levels in tumor and normal samples were also evaluated. Fibrolamellar cancer was confirmed by demonstrating presence (tumor) and absence (normal) of the DNAJ-PKAc oncoprotein (**E**).

We next evaluated for mediators of the cell cycle that could explain the difference in cell cycle progression induced by the presence of *DNAJB1-PRKACA* in the H33 cells. We did not find differences in expression of various cell cycle genes that would explain the phenotype in H33 cells (**Supplemental Data 1 and 2**). For instance, many of the cyclins (*e.g.*, cyclin D1 [*CCND1*] and cyclin E1 [*CCNE1*]) were downregulated as opposed to upregulated, or gene expression levels were nearly undetectable (*e.g*., *CDKN2A* logCPM = -0.8). Because we had identified a more rapid transition from G1 to S and a much greater percentage of cells progressing to G2/M, we postulated that a key regulator of the cell cycle was dysregulated in the *DNAJB1-PRKACA* expressing H33 cells versus HepG2 cells. One such mediator of the cell cycle that modulates activity of several cyclin-dependent kinases (CDKs) is CDK7; CDK7 binds cyclin H and MAT1 to form the CDK-activating kinase (CAK) which phosphorylates target CDKs (CDK1, 2, 4, 6).^26, 27^ We found that CDK7 regulated genes were significantly upregulated in H33 compared to HepG2 cells, including *GATA2*, *GATA3*, *ETV1* and *RUNX1* (all FDR *q* < 0.0001) (**Figure 3B**). Increased expression of the CDK7 regulated genes was confirmed with our separate validated clone (H12 vs HepG2) (**Figure 3B** and **Supplemental Data 3**). Concordantly, the RNA expression data comparing human FLC (n = 35) to normal liver (n = 10) demonstrated elevated expression of *GATA2* (418 [242.8] vs 165 [134], *q* < 0.001), *ETV1* (953 [654] vs 318 [139], *q* < 0.001) and *RUNX1* (3090 [3592] vs 1885 [1367], *q* = 0.018) in the FLC tissue (**Figure 3B**),^25^ which suggested aberrant CDK7 activation in FLC, but clearly required additional confirmation.

While the assessment of certain CDK7 regulated genes (*e.g*., *GATA2*, *ETV1* and *RUNX1*) was an important clue for identifying a putative target downstream of DNAJ-PKAc, a better approach for detection of aberrant CDK7 activation is through phosphoprotein analysis. CDK7 is unique amongst the CDKs in that it is has a dual role as a master regulator of cell cycle (*i.e*., phosphorylates CDK1, CDK2) and a transcriptional regulator (*i.e*., phosphorylates RNA Polymerase II at promoter and enhancer regions) (**Figure 3C**).^26-28^ CDK7 modulates transcription by phosphorylating RNA Polymerase II at specific serine sites in the c-terminal domain (CTD), including serine-5 (preferential target), serine-7 and potentially (much lesser extent) serine-2.^28^ Thus, to complement the human and cell line transcriptional data, we assayed for aberrant activation of CDK7 including phosphorylation of threonine160 on CDK2 (pCDK2) and phosphorylation of RNA Polymerase II (serine-5, serine-7 and serine-2 of C-terminal domain). We performed immunoblotting in the *DNAJB1-PRKACA* expressing H33 cells and parent HepG2 cells and found increased levels of phosphorylated CDK7 and concomitantly increased phosphorylation of CDK2 as well as serine residues in the CTD of RNA Pol II (**Figure 3D**), substantiating the transcriptional data that indicated increased activation of CDK7. To understand whether these findings were relevant in human FLC, we isolated protein from human samples of FLC and paired normal liver and performed an immunoblot assay to assess for CDK2 and RNA Polymerase II (RNA Pol II). We found concordant results with clear increases in serine-5 phosphorylation (sample FCF 57, 82, 83 and 84), serine-7 phosphorylation (sample FCF57, 82 and 83), and increased CDK2 phosphorylation (sample FCF 57 and 84) in the human tumor tissue versus human normal liver (**Figure 3E**). Collectively, this is the first direct evidence of aberrantly increased CDK7 activity in *DNAJB1-PRKACA* expressing cells including phosphorylation of RNA Pol II, phosphorylated (T160) CDK2, and increased expression of CDK7 regulated genes. Most importantly, our cell line data was recapitulated in human samples of FLC, in which RNA and protein analysis confirmed aberrant CDK7 activation. Given the dual role of CDK7 in cell cycle regulation and transcriptional regulation,^26^ this data spurred us to investigate the functional relevance of CDK7 in FLC.

### CDK7 is a novel regulator of FLC-specific gene expression

Using a potent and specific CDK7 inhibitor that is currently in use in clinical trials (SY-5609),^29^ we assessed the efficacy of CDK7 inhibition in the *DNAJB1-PRKACA* expressing H33 cells. We evaluated whether CDK7 modulated the increased phosphorylated CDK2 (cell cycle) and phosphorylated RNA Pol II (transcription) that we had seen in *DNAJB1-PRKACA* expressing H33 cells compared to HepG2 cells. With SY-5609 treatment, we identified clear decreases in phospho-CDK2 and suppressed phosphorylation of serine-2, -5, and -7 (CTD) of RNA Pol II (**Figure 4A**). These data support the idea that CDK7 mediates these specific *DNAJB1-PRKACA* driven changes in H33 cells versus HepG2 cells.

**Figure 4.**
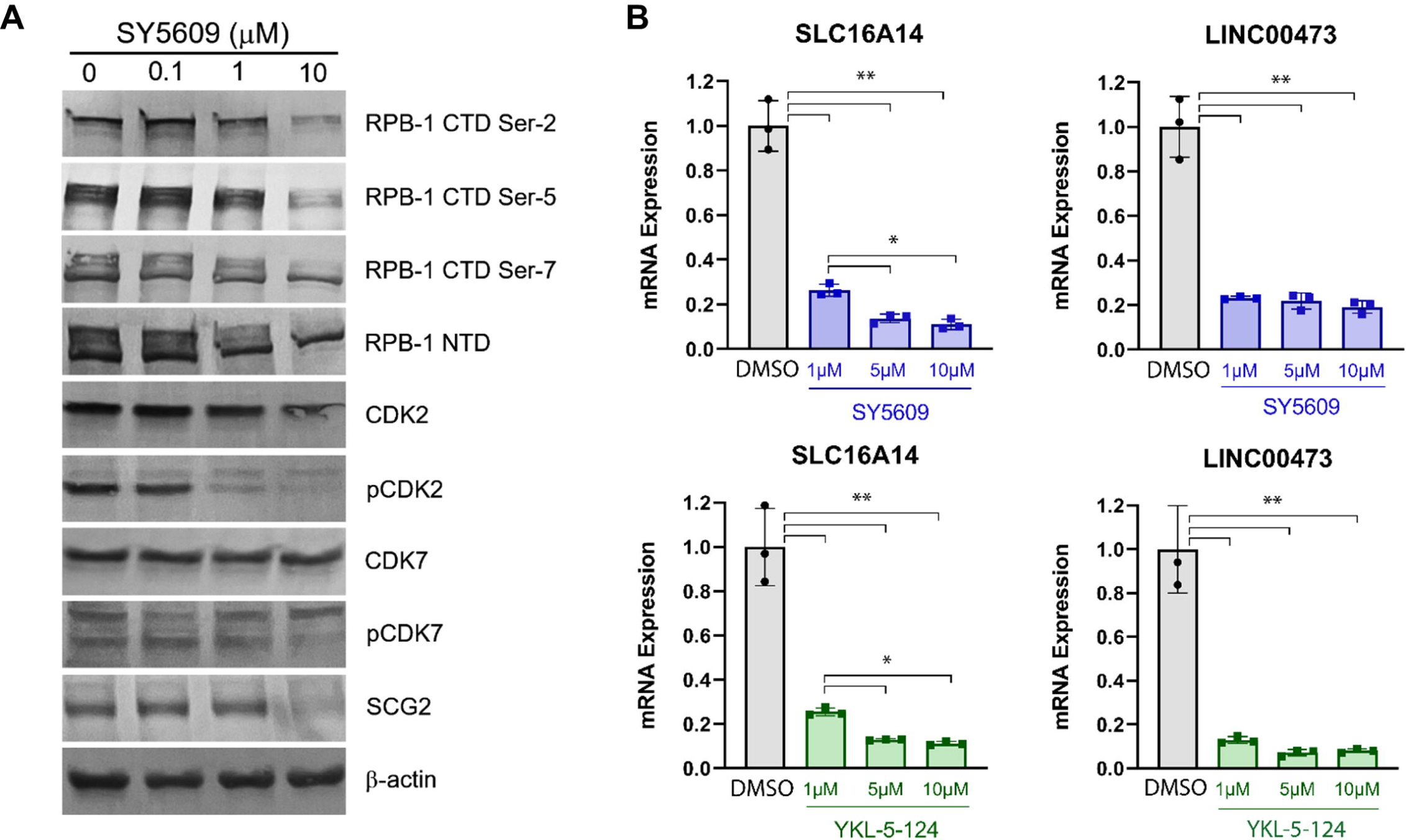
*CDK7 regulates FLC-specific gene expression*. *DNAJB1-PRKACA* expressing H33 cells were treated with a selective CDK7 inhibitor, SY5609 (100 nM, 1 μM, 10 μM), or DMSO control (0) for 24 hours. Known primary targets of CDK7 activation were assessed including RNA Pol II CTD (ser-2, -5, and -7) and Thr160 phosphorylated CDK2 (pCDK2) (**A**). To assess for CDK7-dependent expression of FLC-specific genes, H33 cells were treated with SY5609 (1 μM, 5 μM, 10 μM) and levels of mRNA expression (RT-qPCR) versus DMSO control were evaluated, including *SLC16A14* and *LINC00473*. This was repeated with a separate covalent-binding selective and specific CDK7 inhibitor (YKL-5-124). Shown are three biological replicates per drug dose per mRNA (\*\**p* < 0.0001, \**p* < 0.05) (**B**).

Prior work has shown that in human FLC there is aberrant super enhancer activity due to the DNAJ-PKAc oncoprotein, with consequent increased expression of FLC specific genes including *SLC16A14* and *LINC00473*.^11^ Notably, *SLC16A14* and *LINC00473* are highly expressed in human FLC compared to normal liver and are not elevated in other liver cancers (such as cholangiocarcinoma or HCC).^11^ The function of these genes in FLC and the impact of their marked overexpression (compared to paired normal liver) remain unknown but the products of these genes may serve as unique therapeutic vulnerabilities in FLC and is the focus of current investigations.^11^ RNA Pol II (along with other cofactors and chromatin regulators) is known to be highly enriched at super enhancers – and can transcribe enhancer RNA that may contribute to enhancer function.^30^ Given the importance of CDK7 in facilitating RNA Pol II function (via phosphorylation), and the known role of CDK7 in cancer associated super enhancers,^31-34^ we tested whether the aberrant CDK7 activation may be driving the expression of these FLC-specific super enhancer associated genes.

We treated *DNAJB1-PRKACA* expressing H33 cells with SY-5609 and found a substantial downregulation of both *SLC16A14* and *LINC00473* expression (**Figure 4B**), suggesting a critical role of CDK7 in promoting super enhancer driven gene expression in FLC. To confirm these results, we used a separate potent and selective covalent CDK7 inhibitor (YKL-5-124),^35^ and found identical downregulation of *SLC16A14* and *LINC00473* expression (**Figure 4B**). These data demonstrate for the first time that *SLC16A14* and *LINC00473* are novel CDK7 regulated genes, and that CDK7 – which mediates phosphorylation of RNA Pol II in *DNAJB1-PRKACA* expressing cells – is responsible for regulating expression of FLC-specific super enhancer driven genes. Furthermore, and of substantive clinical importance, CDK7 inhibition can be used to target *both* SLC16A14 and LINC00473 in FLC, which is a key finding given the current inability to inhibit these highly promising targets in FLC (*i.e.*, they are currently not ‘druggable’).

We noted that other putative FLC-specific targets exist that are consistently differentially expressed across multiple FLC datasets,^11, 17^ which matched our H33 vs HepG2 RNA sequencing data. One of prime interest is SCG2, a protein in the neuroendocrine family of secretory proteins. SCG2 is known to be expressed in several tissue types (*e.g.,* adrenal gland, brain, small intestine), but is not typically expressed in liver, which is why the high levels of expression in human FLC compared to normal liver are consequential.^36^ We confirmed our cell line and human database mRNA results with SCG2 protein immunoblotting, which revealed an absence of protein expression in HepG2 cells compared to strong expression in H33 cells (**Figure 3D**). Identical results were seen in human FLC samples versus paired normal liver (**Figure 3E**). SCG2 has been found to promote neuroendocrine differentiation in prostate cancer, a quality that is associated with relapse and treatment-resistance disease.^37^ *SCG2* is presently overexpressed across FLC datasets,^11, 17^ similar to *SLC16A14* and *LINC00473*, and may therefore also be associated with aberrant super enhancer activity. Thus, we postulated that it too could be regulated by CDK7. We treated *DNAJB1-PRKACA* expressing H33 cells with SY-5609 and found that SCG2 expression was dramatically reduced with CDK7 inhibition (**Figure 4A**). We therefore uncovered that a putative target in FLC (SCG2) may be regulated by CDK7 and suppressed by CDK7 inhibition.

### CDK7 inhibition suppresses cancer cell growth in human models of FLC

Given our evidence of CDK7 controlling expression of compelling FLC-specific therapeutic targets (*i.e.*, SLC16A14 and LINC00473),^25^ and the known dual role of CDK7 on cell cycle and RNA Pol II transcriptional regulation,^38, 39^ we hypothesized that the convergent mechanism of CDK7 on cell cycle and super enhancer-driven transcriptional changes would provide a therapeutic vulnerability in FLC cells. We therefore tested whether CDK7 inhibition impacted cell proliferation in H33 cells. SY-5609 demonstrated a strong dose-dependent reduction in cell viability, which was confirmed in a separate clone (H12) (**Figure 5A**). To ensure that the effect we were seeing was CDK7 dependent, we assessed another inhibitor, YKL-5-124 which irreversibly and covalently binds CDK7. YKL-5-124 treatment resulted in a similar dose-dependent reduction of H33 cell viability (**Figure 5A**). As an additional confirmation, we used siRNA against CDK7 and again demonstrated a strong dose-dependent response (**Supplemental Figure 3**), thus revealing that targeting CDK7 dramatically impairs cell viability (*i.e.,* a therapeutic vulnerability) in the FLC model of *DNAJB1-PRKACA* expressing H33 cells.

**Figure 5.**
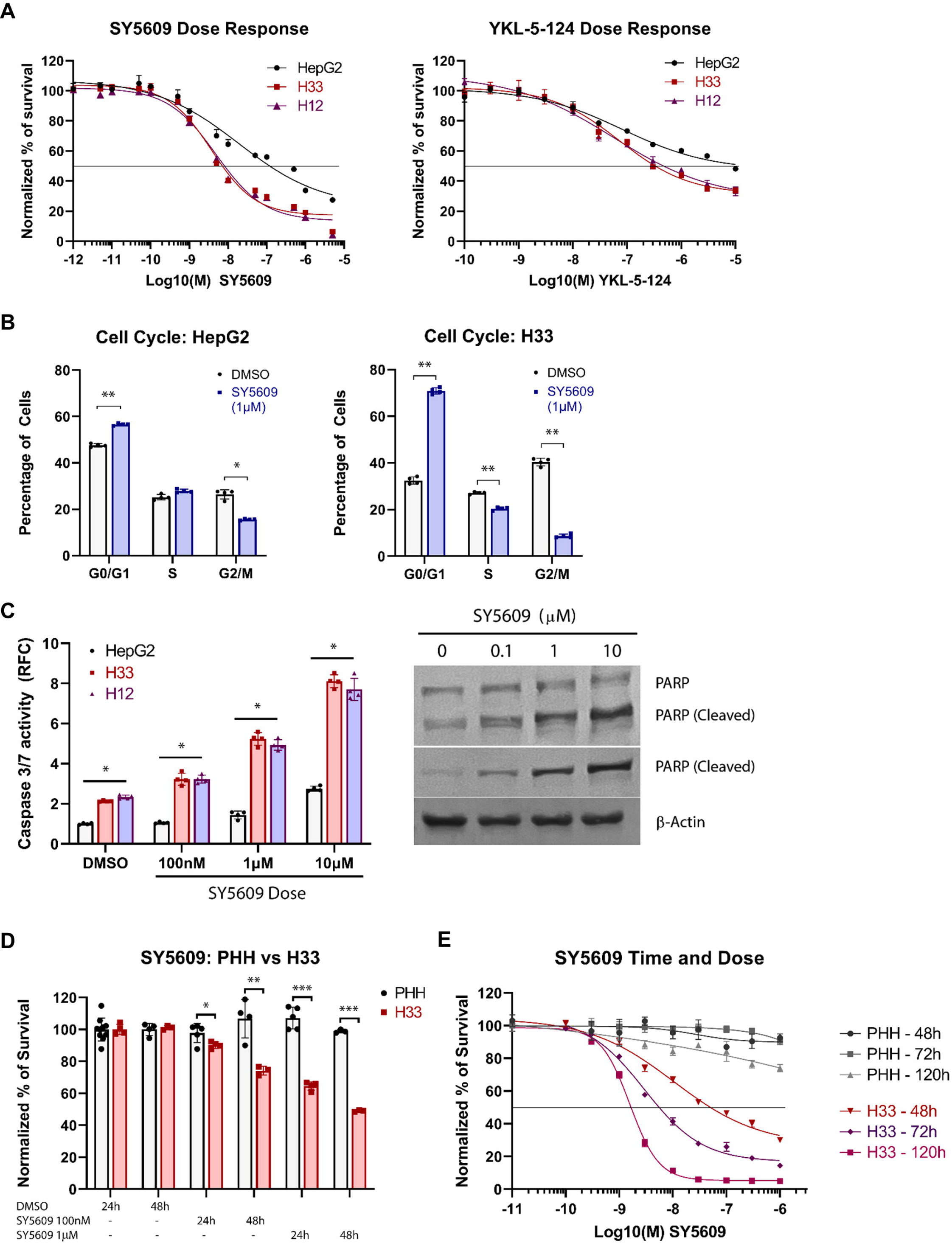
*CDK7 is a novel therapeutic target in DNAJB1-PRKACA expressing cells*. To assess for CDK7 effect on cell viability, HepG2 cells and H33 cells underwent 48hr drug treatment with either SY5609 (1 pM – 5 μM) or YKL-5-124 (100 pM – 10 μM). Percent viability was determined by normalizing to control (DMSO treated). To confirm the findings in the *DNAJB1-PRKACA* expressing H33 cells, a separate clone (H12) was tested with the same drugs over the same dose range. In each figure, the LC_50_ (IC_50_) is represented by the straight line. (**A**). HepG2 cells and H33 cells were synchronized and treated with either DMSO or SY5609 (1 μm) for 24 hours. Percent of cells in G0/G1, S and G2/M were determined by flow cytometry. Shown are four biological replicates per cell line per treatment (\**p* = 0.0015, \*\**p* < 0.0001) (**B**). HepG2 cells and H33 cells were treated with DMSO (control) or increasing doses of SY5609 for 24 hours and Caspase 3/7 activity measured. To confirm the increased apoptotic activity in the H33 cells, a separate clone (H12) was utilized. (\**p* < 0.0001, for H33 vs HepG2 and H12 vs HepG2). To validate the results, PARP and cleaved-PARP (marker for apoptosis) protein were evaluated in HepG2 and H33 cells, using two separate antibodies that either recognize both PARP and cleaved-PARP (top bands) or only cleaved-PARP alone (bottom band) (**C**). Primary human hepatocytes (PHHs) isolated fresh from human donor liver transplant specimens and H33 cells were treated with DMSO, SY5609 100 nM, or SY5609 1 μM for 24- and 48-hours. The percent of viable cells was determined by normalizing to DMSO control. There were four biological replicates per group per time/treatment dose (\**p* = 0.02, \*\**p* = 0.01, \*\*\**p* < 0.0001) (**D**). PHHs and H33 cells were treated with SY5609 (100 pM – 5 μM) for 48-, 72- or 120-hours and percent viability assessed, normalized to DMSO control. The LC_50_ (IC_50_) is demarcated by the solid line. There were four biological replicates for each drug dose at each time point for each line (PHH and H33) (**E**).

The marked reduction in cancer cell viability due to CDK7 inhibition may be due to cell cycle arrest, induction of apoptosis, or both. Due to the significant differences in cell cycle progression between H33 cells and parent HepG2 cells, we first measured the effect of CDK7 inhibition on cell cycle. We found that SY-5609 treatment resulted in a modest effect in the HepG2 cells, with a moderately reduced percentage of cells in G2/M (26.4% DMSO vs 15.5% SY-5609, *p* = 0.0015) and a reciprocal increase in the percent of cells in G0/G1 (47.6% DMSO vs 56.6% SY-5609, *p* < 0.0001). In contrast, CDK7 inhibition in H33 cells resulted in a profound cell cycle shift with most cells in G0/G1 (32.5% DMSO vs 70.9% SY-5609, *p* < 0.0001) and a marked reduction in the percent of cells in G2/M (40.4% DMSO vs 8.7% SY-5609, *p* < 0.0001) (**Figure 5B**). This apparent arrest in G0/G1 due to CDK7 inhibition reflects the integral nature of CDK7 in promoting cell cycle progression in *DNAJB1-PRKACA* expressing cells. We subsequently examined the effect on apoptosis with CDK7 inhibition and expected that the dominant effect would be cell-cycle mediated. However, SY-5609 treatment of H33 cells demonstrated a significant increase in caspase 3/7 activity compared to HepG2 cells at all treatment doses (SY-5609 100 nM, 1 μM, 10 μM, all *p* < 0.0001), which was confirmed with the separate H12 clone, demonstrating high sensitivity of *DNAJB1-PRKACA* expressing cells to CDK7 inhibition. We again validated the significant increase in apoptosis through detection of cleaved PARP1, which displayed a clear dose-dependent increase in cleaved protein with SY-5609 treatment (**Figure 5C**). Together, this data shows that CDK7 is a critical component of cell proliferation / viability in *DNAJB1-PRKACA* expressing cells, and likewise, CDK7 may represent a key therapeutic vulnerability to exploit in FLC.

A necessary component of any potential therapy is the identification of a true therapeutic window. Because toxicity may limit translational potential (*i.e.,* therapy tested must be tolerated), we assessed the effect of CDK7 inhibition in human liver cells. We isolated human hepatocytes immediately from fresh liver (transplant donor specimen) and examined the impact of SY-5609 at two separate doses (100 nM and 1 μM) for 24- and 48-hours (**Figure 5D**). While the H33 cells demonstrated clear dose and time-dependent sensitivity to SY-5609 treatment, there was no change in cell viability in the primary human hepatocyte (PHH) liver cells compared to DMSO control, confirming the lack of toxicity to normal liver cells at the doses tested. We confirmed our findings with separate dose-response experiments in the human hepatocytes at 48-, 72-, and 120-hours treatment with SY-5609 (**Figure 5E**), to ensure minimal toxicity even with increased duration of exposure. Notably, with increasing time, the LC_50_ (IC_50_) value of SY-5609 decreased in the H33 cells (IC_50_ shifts to the left: 7.97E-8, 8.67E-9 and 1.71E-9 at 48-, 72- and 120-hours respectively), indicating both a dose and time-dependent cell death. Meanwhile, there was minimal impact on the human hepatocytes, even out to 120-hours treatment duration. We therefore identified a therapeutic vulnerability in *DNAJB1-PRKACA* expressing H33 cells that was not present in the human fresh liver tissue (*i.e*., no cytotoxicity), indicating an excellent therapeutic window/index for CDK7 inhibition.

To better understand the clinical relevance of our results, we tested CDK7 inhibition independently in five separate patient-derived models from two separate laboratories. Using a patient-derived FLC cell line that has been well characterized (FLC-H),^10, 11^ we found a suppression of cell growth after SY-5609 treatment compared to control treated cells, with a greater than 50% reduction in cell viability (SY-5609 1 μM, *p* < 0.01) (**Figure 6A**). Using two distinct patient-derived FLC cell lines (FLC1025 derived from primary tumor, FLCMet derived from metastatic disease) from a separate laboratory, we identified exquisite sensitivity to SY-5609 treatment with a dose-dependent reduction in cell viability, confirming the findings from our H33 cells and the FLC-H cell line (**Figure 6B**). We next tested the effect of SY-5609 in tissue slices derived from human FLC (**Figure 6C**), and from a separate set of tissue slices derived from a patient-derived xenograft of FLC (**Figure 6D**). Remarkably, we found a similar reduction in cell viability compared to the patient-derived cell lines, which confirmed the efficacy of CDK7 inhibition (*i.e.*, validated in 5 separate patient-derived model systems). Thus, we have uncovered a novel druggable pathway in FLC that demonstrates aberrantly heightened activity due to DNAJ-PKAc (*i.e.,* in FLC vs normal liver and in H33/H12 vs HepG2) and appears to be a central regulator of known FLC-specific targets (**Figure 7**). This link between the fusion oncoprotein and downstream super enhancer driven genes represents a key therapeutic vulnerability. Consequently, inhibition of this pathway could be combined with another novel therapy (*e.g*., small molecules targeting DNAJ-PKAc) as a powerful combination strategy. As proof of principle, CDK7 inhibition is known to be effective in cancers with low levels of expression of anti-apoptosis proteins.^40^ We reasoned that CDK7 would then likely synergize with anti-apoptosis inhibitors and screened SY-5609 with various anti-apoptosis inhibitors. Interestingly, while there was low toxicity of the combination therapy in fresh human liver tissue (PHHs), SY-5609 showed strong synergy with the BCL-xL inhibitor A1331852 (**Figure 6E, Supplemental Figure 4**).^41^ This was shown at very low doses of A1331852 (10 nM), which is an important consideration due to the known toxicity profile of anti-apoptosis inhibitors in humans with overlapping risks for myelosuppression. This is also notable due to the recent seminal work demonstrating the synergistic value of BCL-xL inhibitors in FLC.^17, 42^ Additional combination screening will need to be performed to identify the top synergistic candidates with CDK7 inhibition, to optimize the tumoricidal effect while maintaining low toxicity. This will be a focus of future efforts.

**Figure 6.**
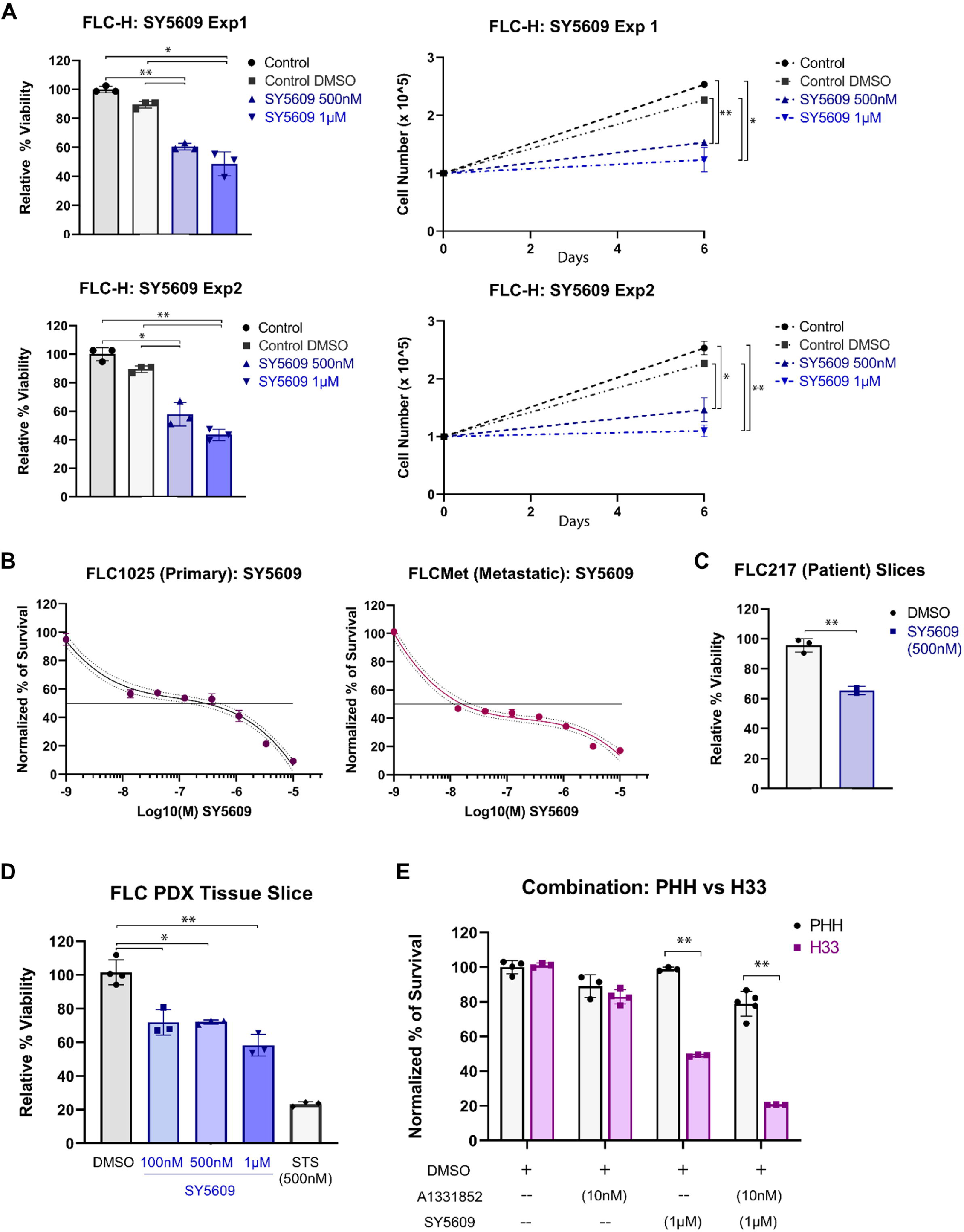
*CDK7 inhibition is lethal to human FLC*. An FLC cell line derived from human FLC (FLC-H) was grown for six days (control) or treated with DMSO (Control DMSO) versus SY5609 at two separate doses (500 nM and 1 μM). Survived cells were quantified at day zero and day six. Additionally, the day six percent survival compared to controls was calculated. Shown are two separate experiments with three biological replicates per experiment (\**p* < 0.01, \*\**p* < 0.001 versus Control and Control DMSO) (**A**). In a separate independent laboratory, an FLC cell line was derived from a patient’s FLC liver tumor (FLC1025), and a separate cell line was derived from another patient’s metastatic FLC tumor implants (FLCMet). Each line was treated with SY5609 (10 nM – 10 μM). Percent survival was determined normalized to DMSO control, with three biological replicates per dose. Shown is dose response with curve (and 95% CI) and LC_50_ (IC_50_) demonstrated by the straight line (LC_50_ FLC1025 ∼ 300 nM, LC_50_ FLCMet ∼ 20 nM) (**B**). Human tissue slices derived from a patient with FLC (FLC217) were treated with DMSO or SY5609 (500 nM), and percent viability determined compared to DMSO control (\*\**p* = 0.003) (**C**). In a separate experiment, tissue slices derived from human FLC grown in a patient derived xenograft (PDX) model were treated with DMSO, SY5609 (100 nM, 500 nM and 1 μM) and Staurosporine (STS) 500 nM (positive control). There were 3-4 biological replicates per group (\**p* < 0.01, \*\**p* < 0.001) (**D**). Primary human hepatocytes (PHH) and H33 cells were treated for 48 hours with a selective BcL-xL inhibitor (A1338152, 10nM dose), SY5609 (1μM) or combination A1331852 (10nM) + SY5609 (1μM), and percent survival in each group determined by normalizing to DMSO control. There were 3-5 biological replicates for each treatment in PHH and H33 cells (\*\**p* < 0.001) (**E**).

**Figure 7.**
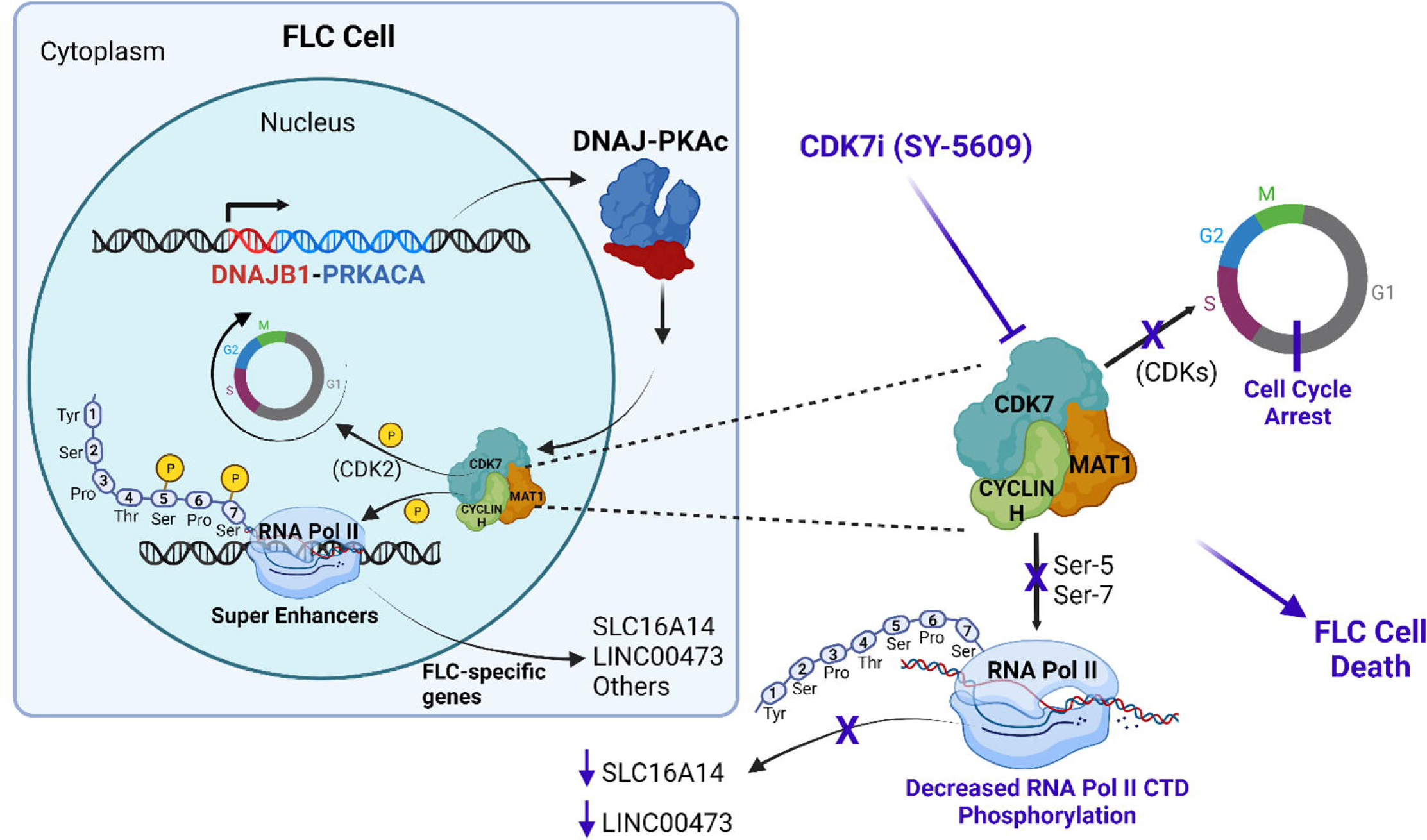
*Aberrant CDK7 activity is a therapeutic target in DNAJ-PKAc expressing cells*. The *DNAJB1-PRKACA* oncogene fusion is the driver mutation in nearly all human FLC, and the protein product (DNAJ-PKAC) causes aberrant activation of downstream targets through dysregulated kinase activity. Consequently, the DNAJ-PKAc oncoprotein results in reprogramming of enhancer regions to generate high levels of expression of FLC specific genes, such as *SLC16A14* and *LINC00473*, and drives tumor cell growth / metastasis. It has previously been indeterminate how DNAJ-PKAC is linked to the emergence of super enhancers and transcriptional changes. This study uncovers aberrant CDK7 activation in *DNAJB1-PRKACA* expressing cells, resulting in more rapid cell cycle progression and mediation of FLC-specific gene expression likely through RNA Pol II phosphorylation. While it is uncertain how DNAJ-PKAc promotes dysregulated CDK7 activity (*i.e.*, direct interaction or through intermediate protein signaling), it does appear that CDK7 is a key regulator of cell proliferation in *DNAJB1-PRKACA* expressing cells. In turn, CDK7 inhibition (e.g., SY-5609) causes a robust cell cycle arrest, decreased RNA Pol II CTD phosphorylation, and suppressed FLC-specific gene expression, and targeting CDK7 represents a therapeutic vulnerability in several different models of human FLC.

## DISCUSSION

Since its initial discovery, there has been inconclusive evidence of how DNAJ-PKAc drives cellular changes to promote neoplastic transformation and progression of FLC. Concordantly, we developed a systematic approach to identify and subsequently validate a potential target. We first introduced the *DNAJB1-PRKACA* gene fusion at the native promoter of HepG2 cells to investigate pathophysiologic mechanisms associated with the oncogene fusion. After rigorous validation of the generated *DNAJB1-PRKACA* expressing cell line, we used it as a tool to uncover a key component of DNAJ-PKAc driven pathogenesis: aberrant CDK7 activation. We confirmed our discovery of abnormally increased CDK7 activation in human tumor vs normal tissue and human-derived cancer cells. Importantly, there was strong concordance of results between our *DNAJB1-PRKACA* expressing cells (vs parent cell line), the human tumor samples (vs normal liver), and among the five separate human FLC-derived models. These findings were consistent across three independent laboratories, underscoring the clinical relevance of our results. Thus, CDK7 appears to qualify as a central regulator in FLC, such that *DNAJB1-PRKACA* expressing cells (but not control cells) possess aberrant activation of CDK7, and the heightened CDK7 activity drives expression of super enhancer driven genes in FLC.

The research landscape in FLC changed after a 2014 pioneering study that revealed the presence of the *DNAJB1-PRKACA* gene fusion in human FLC, and was later shown to drive FLC in mouse.^5, 10, 11^ The clinical importance of *DNAJB1-PRKACA* in human FLC was subsequently confirmed in two separate studies, including one in which the authors utilized shRNA spanning different regions of the *DNAJB1-PRKACA* breakpoint to demonstrate that the FLC cells were reliant upon *DNAJB1-PRKACA* for cell survival (oncogenic addiction).^12^ This indicates that following tumor initiation, FLC requires the presence of DNAJ-PKAc (and downstream signaling) to survive. Separately, a group with expertise in vaccine therapy and immune targeting of cancer exploited the fact that the *DNAJB1-PRKACA* fusion transcript generates a protein product (DNAJ-PKAc) that is the clonal oncogenic driver in FLC, and consequently, FLC displays tumor-specific neoantigens that can be recognized by the immune system.^13^ The authors found that unique fusion neoepitopes were predicted as ligands to the majority of HLA class II alleles and of HLA class I allotypes, and that CD8^+^ T-cells would recognize the neoepitopes, which was confirmed *in vitro*. They constructed a vaccine comprised of three separate HLA class I ligands (9-10 amino acids in length) and one longer peptide predicted to bind HLA class II (DP) to provide a personalized vaccine for a patient who had undergone multiple lines of surgery (including transplant) and systemic therapy. The vaccine was administered with a toll-like receptor 1/2 agonist, and the patient developed neoepitope-specific T-cells with objectively high intensity response to *DNAJB1-PRKACA* derived neoepitopes *ex-vivo*, as well as durable clinical response (no disease relapse at 21 months post-vaccination). Presumably, cells expressing DNAJ-PKAc were identified and targeted, and the absence of recurrence indicates that cancer cells could not ‘escape’ by selection for cancer cells without *DNAJB1-PRKACA*. Thus, combined, these two studies highlight the critical importance of *DNAJB1-PRKACA* in FLC, and that this fusion uniquely promotes a pathogenic mechanism within the cancer cells that is specifically reliant upon the presence of the DNAJ-PKAc protein.

Therefore, identifying how DNAJ-PKAc modulates downstream targets to drive cell growth and metastasis should similarly reveal a critical molecule or set of molecules that relay the aberrant signal. Because evidence from a recent comprehensive analysis of human FLC revealed downstream reprogramming of enhancer regions to drive high levels of transcription of FLC-specific genes,^11^ the critical molecule or set of molecules should similarly impact these FLC-specific genes. Conceptually, this then provides two separate ‘set-points’ in human FLC, where irrefutable evidence demonstrates the upstream driver of cellular changes (*i.e*., DNAJ-PKAc oncoprotein), and the downstream output as a consequence of neoplastic change (*i.e*., FLC-specific gene expression including *SLC16A14* and *LINC00473*).^6, 11-13^ Our target (CDK7) links these two ‘set-points’, where the introduction of *DNAJB1-PRKACA* promotes aberrant activity of CDK7, and suppression of CDK7 inhibits expression of FLC-specific genes.

CDK7 is a serine/threonine protein kinase that exists as a CDK-activating kinase (CAK) and has gained significant interest as a target in cancer due to its dual roles in cell cycle regulation and transcriptional regulation.^39, 43^ CDK7 targets the heptad sequence (Tyr-Ser-Pro-Thr-<u>Ser</u>-Pro-Ser) of the CTD in RNA Pol II, with preferential phosphorylation of serine-5 in the sequence.^38^ This is notable due to the fact that among the serine targets, we identified serine-5 as the most differentially phosphorylated in H33 versus HepG2 cells and more importantly, in human FLC tumor versus normal liver. Meanwhile, CDK substrates (*e.g*., CDK1, 2) are phosphorylated on the threonine residue (<u>Thr</u>-X-X-Val-Val-Thr-Leu), with Thr160 as the phospho-acceptor in CDK2; this increased pCDK2 (Thr160) is what we found in our H33 samples and some human FLC samples. The location of CDK2 phosphorylation is interesting due to the preference of CDK7 to phosphorylate serine or threonine residues with an adjacent proline (+1 position).^38^ Although the target substrates are structurally unrelated (*e.g*., RNA-Pol II CTD and CDK2), the ability of CDK7 to precisely and selectively recognize these substrate classes for phosphorylation was disentangled by Larochelle and colleagues.^38^ They found that CDK7 can exist as a dimer (with Cyclin H) or a trimeric complex (CDK7 / Cyclin H / MAT1), and that only the MAT1 bound trimeric complex can phosphorylate RNA-Pol II CTD, the efficiency of which is increased by CDK7 phosphorylation. Meanwhile, both the dimeric and trimeric complex can phosphorylate the various CDK proteins (*e.g*., CDK1, CDK2), which was consistent with results from another study.^43^ Therefore, the distinct mechanisms of CDK7 function result in high specificity and selectivity for substrates, with CDK2 phosphorylation and RNA Pol II CTD phosphorylation representing the divergent outputs of CDK7 activation. This mechanistic insight not only accounts for our phosphoprotein results (*i.e*., preferential serine-5 of RNA Pol II CTD phosphorylation), but also explains the other phenotypic changes that occurred with CDK7 inhibition (*i.e.,* G0/G1 cell cycle arrest, suppression of FLC-specific super enhancer driven gene expression, potent FLC cell death in human models). Of substantial interest, however, is how CDK7 becomes dysregulated in DNAJ-PKAc expressing cells (discussed below).

We utilized selective CDK7 inhibitors to complement our mechanistic analysis and provide insight on the clinical relevance of targeting the aberrantly activated CDK7 pathway. For instance, we used SY-5609 throughout this study because it is a potent and selective CDK7 inhibitor with demonstrated effects on cell cycle, transcription and apoptosis *in vitro* and *in vivo*.^29^ Notably, SY-5609 is currently in use in clinical trials, with a defined safety profile in humans (NCT04247126, NCT04929223).^29^ To confirm that the effects of SY-5609 were CDK7 dependent, we also utilized a separate covalent binding CDK7 inhibitor (YKL-5-124) as well as siRNA to CDK7. Our approach of using multiple methods was informed by a recent study where investigators who developed an older generation CDK7 inhibitor (THZ1) recognized that the effects initially attributed to CDK7 inhibition were instead due to CDK12/13 inhibition *in addition* to CDK7 inhibition.^33, 35^ These investigators then set out to develop a more specific inhibitor that maintained the irreversible binding to Cys312 of CDK7 (mechanism of THZ1). They generated YKL-5-124 by hybridizing the covalent warhead of THZ1 with the pyrrolidinopyrazole group from PF-3758309 (a PAK4 inhibitor) to substantially improve CDK7 selectivity.^35^ Meanwhile, the investigators that developed SY-5609 achieved high potency and selectivity for CDK7 without requiring a covalent warhead, and in an orally bioavailable form.^29^ Through a series of structural modifications of a prior compound (SY-1365 – derived from THZ1), they were able to generate SY-5609, which demonstrated a selectivity window of 4340-fold over the closest off-target (CDK16). In contrast to SY-1365, YKL-5-124 and THZ1, SY-5609 does not covalently bind to CDK7, yet maintains a high specificity through non-covalent interactions. Therefore, we used these two separate compounds as well as siRNA to CDK7 in our initial studies (H33 cells) to support the involvement of aberrant CDK7 function in FLC and confirmed the efficacy of our selected compound (SY-5609) in five separate human models of FLC. Given the substantive data on selectivity for CDK7 of the compounds used, and the observations made in our cell line and human samples, we provide convincing evidence that suppression of CDK7 activity is a novel therapeutic vulnerability in FLC.

There is, however, uncertainty as to whether the impact on cell viability due to CDK7-inhibition was a direct consequence of its role in cell cycle (*i.e.*, CDK2 mediated), or due to its function in RNA Pol II activity (or both). For instance, while the effect of CDK7 inhibition on cell cycle in the *DNAJB1-PRKACA* expressing H33 cells was profound (and could certainly explain the dose-dependent reduction in survival in FLC), CDK7 inhibition was also shown to markedly suppress expression of *SLC16A14* and *LINC00473*. These genes were identified from prior work that established unique super enhancer driven genes in human FLC.^11^ Super enhancers within cancer cells are responsible for high levels of expression of genes that are essential for cell identity and survival.^30, 31^ These key genes associated with super enhancers control cell state and cancer hallmark qualities,^30^ highlighting the critical importance of prior work that identified genes associated with enhancer hotspots (*e.g., LINC00473*).^11^ Concordantly, SLC16A14 and LINC00473 remain highly intriguing and promising targets, which highlights the novel and remarkable finding in this study that CDK7 appears to function as a key regulator of these FLC-specific genes. Further, we have discovered that CDK7 may function as a master regulator of the super enhancer-driven landscape in FLC, similar to prior preclinical data supporting CDK7 as a lead target for super-enhancer-driven cancer types (*i.e.*, MYC).^31, 33, 34^ Here, we provide convincing evidence of how DNAJ-PKAc may impact expression of certain FLC-specific genes through CDK7 activation. Future investigation will focus on elaborating the CDK7-driven effects including differentiating the contribution of cell cycle (*e.g*., CDK2 pathway) and transcriptional machinery (*i.e*., RNA Pol II) such that additional putative targets downstream of CDK7 can be identified for synergistic combination therapy.

Despite the compelling data from this current work, a key question that remains is how DNAJ-PKAc causes aberrant activation of CDK7 to promote cell cycle activity and transcriptional dysregulation. While our discovery is a significant advancement in the mechanistic understanding of DNAJ-PKAc in FLC, the proximity of CDK7 to DNAJ-PKAc is indeterminate. Specifically, it is unknown whether CDK7 is directly phosphorylated by the fusion oncoprotein, or whether intermediate proteins transmit the signal from DNAJ-PKAc. Although there is evidence that native PKA and CDK7 may have overlapping targets (*e.g.,* nuclear retinoic acid receptor alpha),^44^ currently, there is no evidence that native PKA directly regulates CDK7 activity. And while it is known that DNAJ-PKAc induces abnormal cAMP signaling and phosphorylates targets distinct from native PKA,^18, 19^ our findings are the first evidence (to our knowledge) of altered CDK7 pathway activity due to the DNAJ-PKAc oncoprotein. Due to the complexity of CDK7 function (*i.e*., regulation of cell cycle and RNA Pol II is mediated by dimeric vs trimeric complex and phosphorylation status of CDK7),^38, 43^ there is likely a multifaceted interplay between DNAJ-PKAc and CDK7. In future work, we will seek to understand how DNAJ-PKAc modulates CDK7 protein to promote cell cycle changes and transcriptional remodeling.

The data presented here are consequential and support a role of CDK7 in FLC. Targeting of CDK7 induces cell cycle arrest, apoptosis, and suppression of key FLC marker genes. Notably, this work also begins to untangle the regulation of super enhancer driven genes (which are top therapeutic candidates) that are uniquely expressed in FLC as a result of *DNAJB1-PRKACA*.^11^ Collectively, we provide the basis for CDK7 inhibition as a strong candidate for *further investigation* in human FLC.

## Acknowledgements

We would like to acknowledge the University of Wisconsin Biotechnology Center Gene Expression Center and Bioinformatics Resource Center for their contributions to this work. The authors thank the University of Wisconsin Carbone Cancer Center Flow Cytometry Laboratory, supported by P30 CA014520, for use of its facilities and services. The research reported in this publication was supported by a Department of Defense Award [W81XWH-19-1-00546] (TSG), Department of Defense Rare Cancer Research Program Idea Award [RA210165] (SRK), by the American College of Surgeons Faculty Research Fellowship Award (SRK), and the Fibrolamellar Cancer Foundation (SRK, TSG, PS). The content is solely the responsibility of the authors and does not necessarily represent the official views of the Department of Defense.

## Author Contributions

Conceptualization, S.M.R-K. and C.A.B.; Methodology, M.N., S.M.R-K., A.Y., T.S.G. and P.S.; Formal Analysis, M.E.B, S.M.R-K. and A.Y.; Investigation, M.N., C.C., K.Z., I.Y., N.A.K., A.R.F. and M.C.; Resources, S.M.R-K., C.A.B., T.S.G., P.S., A.Y., D.P.A-A and J.D.K.; Data Curation, S.M.R-K.; Writing – Original Draft, S.M.R-K.; Writing – Review and Editing, S.M.R-K., M.N., C.A.B., A.Y., J.D.K., D.P.A-A., T.S.G. and P.S.; Supervision, S.M.R-K., T.S.G and P.S.; Funding Acquisition, S.M.R-K., T.S.G and P.S.

## Declaration of Interests

The authors declare no competing interests.

## Materials Availability Statement

The modified HepG2 cell lines (H33 and H12 cells) created for use in this manuscript are available upon reasonable request from the corresponding author, Sean Ronnekleiv-Kelly.

## Supplemental Data

**Supplemental Figure 1.**
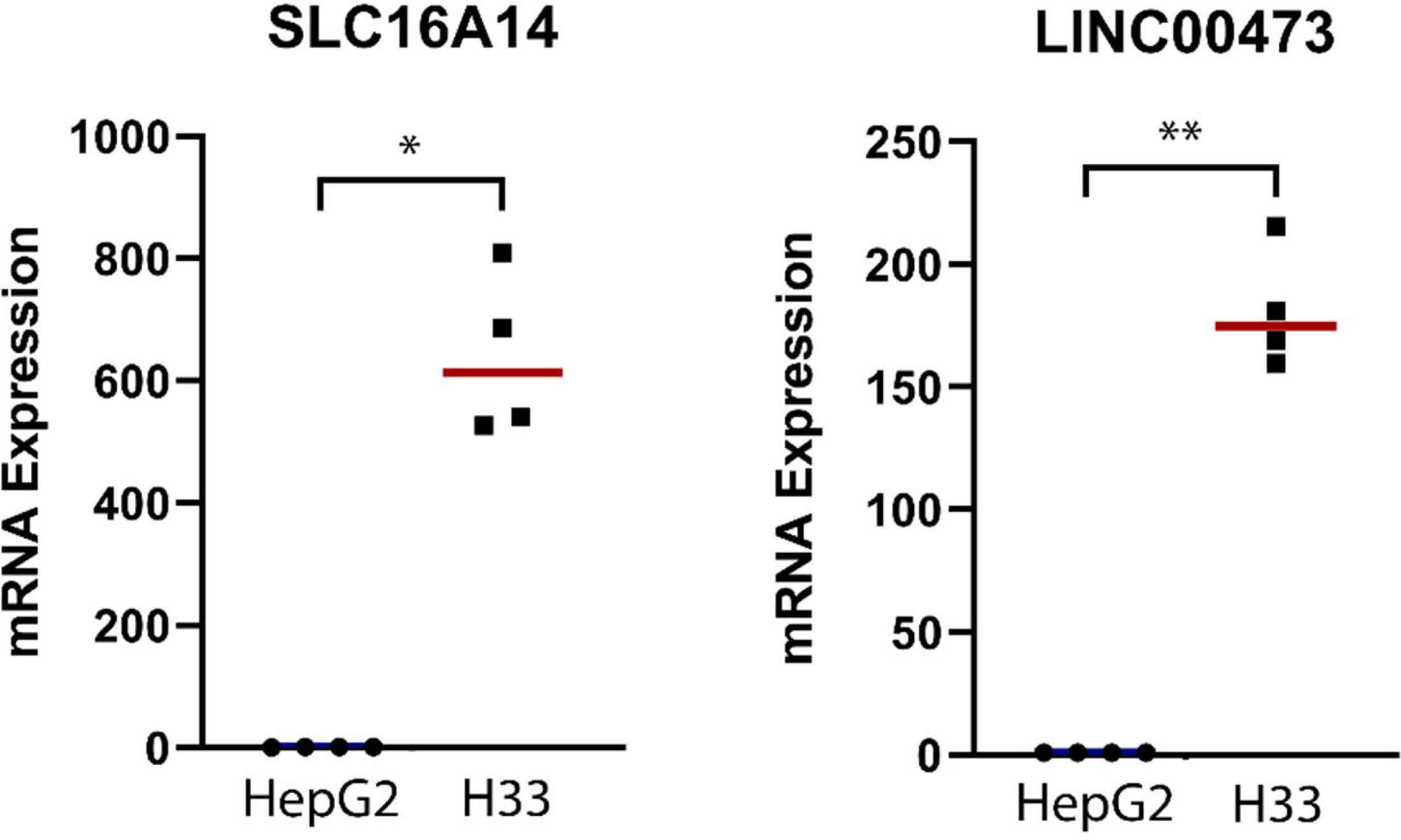
Related to Figure 3. *Quantitative PCR (qPCR) to determine FLC-specific gene expression in H33 and HepG2 cells*. *LINC00473* and *SLC16A14* mRNA expression levels were assessed in the *DNAJB1-PRKACA* expressing H33 cells compared to parent HepG2 cells. Shown is mRNA expression relative fold change compared to HepG2, with four biological replicates per cell line (\**p* = 0.002, \*\**p* = 0.0007).

**Supplemental Figure 2.**
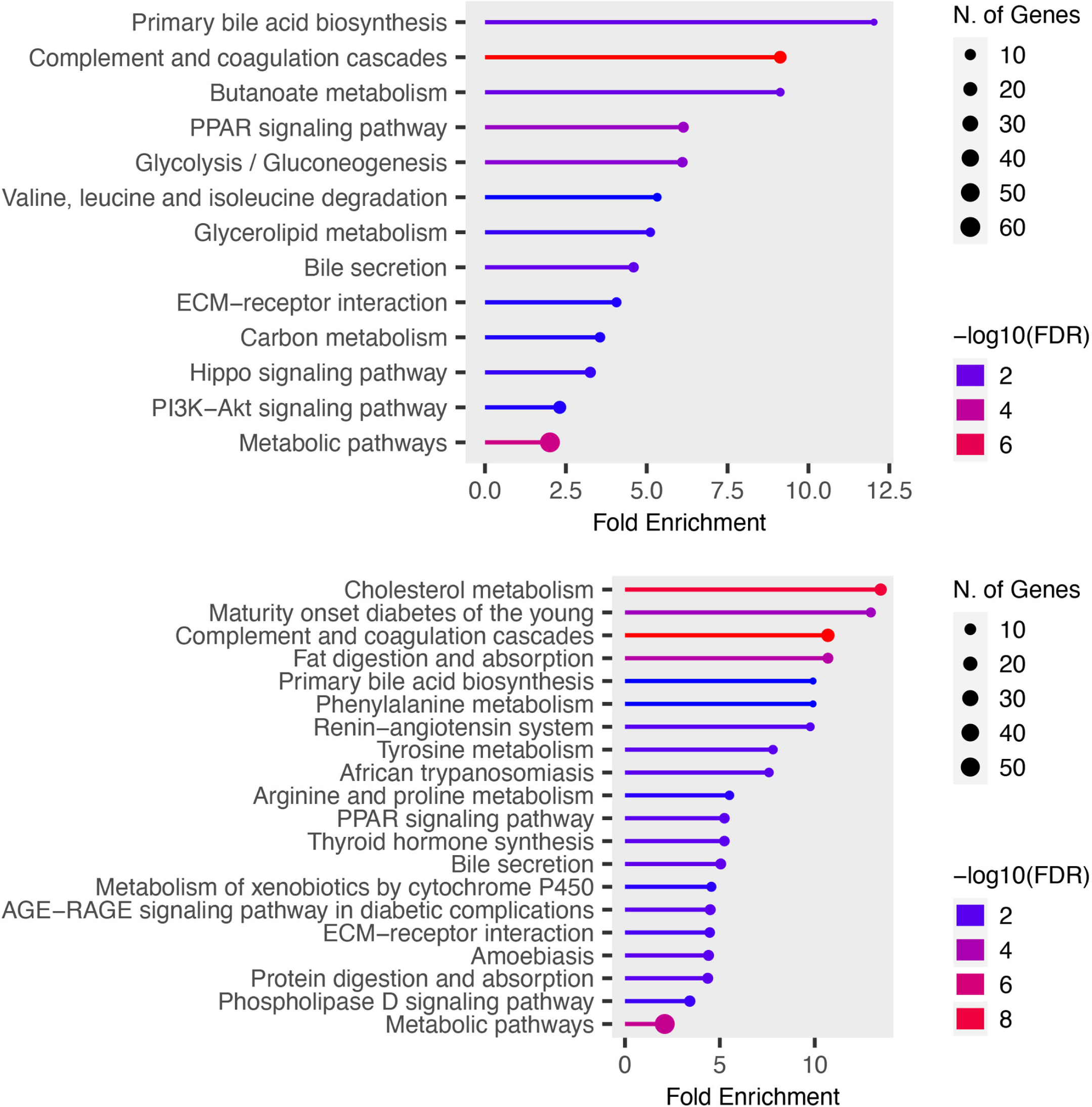
Related to Figure 3. *KEGG pathway analysis*. The top 500 differentially expressed genes in the human FLC (n = 35) versus human normal liver (n = 10) [top figure], and the top 500 differentially expressed genes between H33 cells (n = 6) versus HepG2 cells (n = 6) [bottom figure] were included in the KEGG pathway analysis. Of the thirteen enriched pathways in the human samples, six overlapped with the cell line enriched pathways including i) primary bile acid biosynthesis, ii) complement and coagulation cascades, iii) PPAR signaling pathway, iv) bile secretion, v) ECM-receptor interaction and vi) metabolic pathways. Fold enrichment is shown on the x-axis with specific pathway on the left y-axis. The number of genes involved in each pathway is depicted by circle size and the minus log-base-10 false discovery rate (FDR) is depicted by color.

**Supplemental Figure 3.**
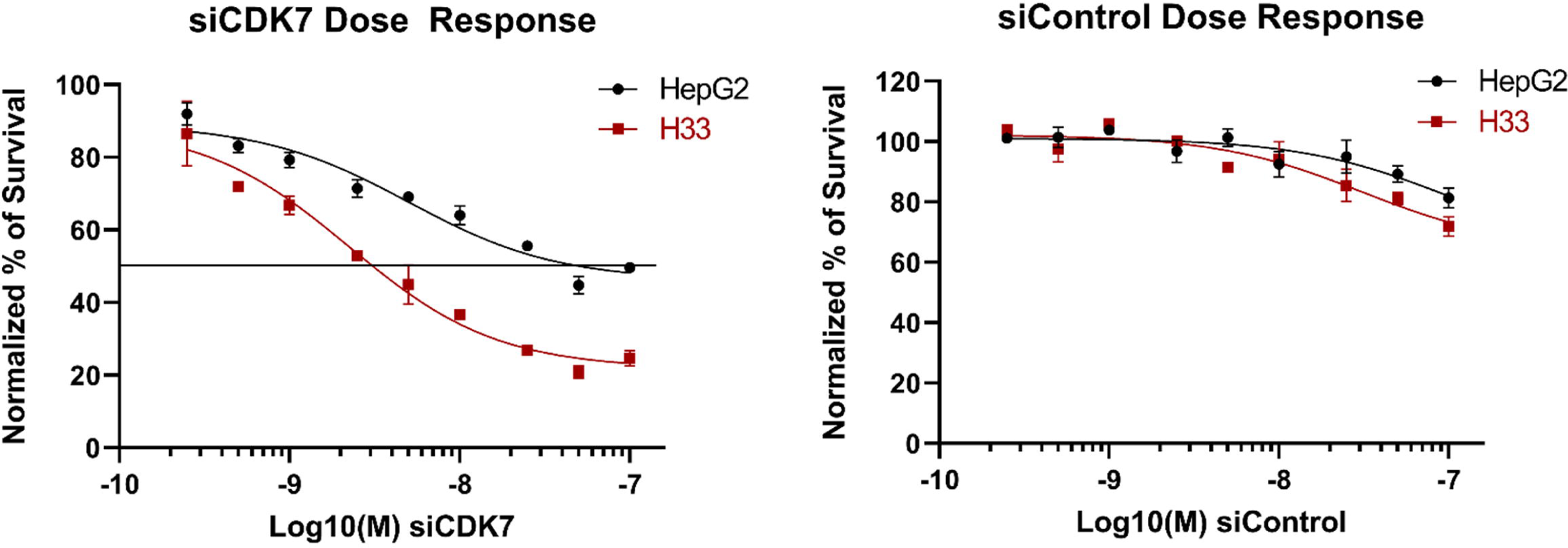
Related to Figure 5. *siRNA dose response in HepG2 and H33 cell lines*. HepG2 cells and H33 cells were treated with siCDK7 (300 pM – 100 nM) or siControl (300 pM – 100 nM). Percent viability was determined by normalizing to control (DMSO) treated cells. The LC_50_ (IC_50_) value is demarcated by the solid black line in the siCDK7 treated cells (IC_50_ ∼3 nM H33 cells vs ∼50 nM HepG2 cells).

**Supplemental Figure 4.**
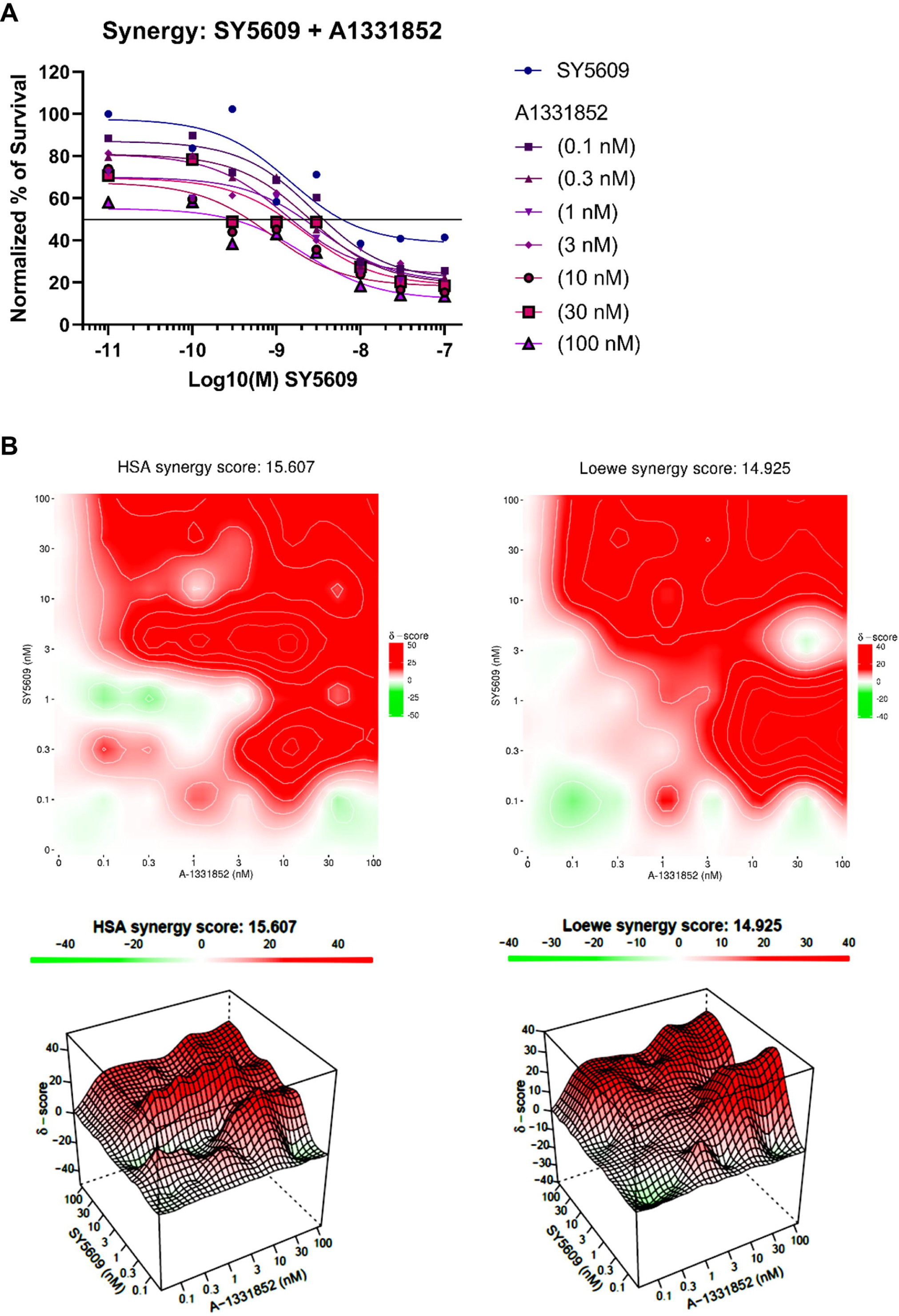
Related to Figure 6. *Synergistic combination therapy SY5609 and A1331852*. H33 cells were treated with combination SY5609 (0 – 100 nM) and A1331852 (0 – 100 nM). Shown is the dose response curve for SY5609 alone (blue) and the same dose response of SY5609 with increasing doses of A1331852. The LC_50_ (IC_50_) value is demarcated by the straight line, demonstrating a shift to the left with increasing concentration of A1331852 (**A**). Synergy was evaluated using the highest single agent (HSA) and Loewe methods (antagonistic < -10, additive -10 < x < 10, synergistic > 10). Depicted is the overall synergy score (15.607 and 14.925, respectively) and heat map representing individual treatment combination scores in 2-dimensional and 3-dimensional maps (**B**).

**Supplemental Data 1**. Related to Figure 3. *Expression table of cell cycle genes in HepG2, H33 and H12 cells*. This table was derived from Supplemental Data 2 and 3.

**Supplemental Data 2**. Related to Figure 3. *Differential expression of HepG2 and H33 cells*. Deposited in GEO.

**Supplemental Data 3**. Related to Figure 3. *Differential expression of HepG2 and H12 cells*. Deposited in GEO.

## METHODS

### Cell line and culture

Hepatoblastoma HepG2 cell line was obtained from ATCC (Manassas, VA). Cells were maintained in RPMI1640 culture medium (RPMI1640 medium (Life Technologies, Carlsbad, CA) supplemented with 10% fetal bovine serum (Gemini Bio Products, Sacramento, CA) and 1% penicillin–streptomycin-glutamine (Life Technologies) at 37 °C and 5% CO2.

### Generation of *DNAJB1-PRKACA* expressing cells

To generate HepG2^DNAJB1-PRKACA^ cells, mammalian dual-guide RNAs-CRISPR-CAS9-EGFP vector was designed (VectorBuilder, Chicago, IL). The guide RNA sequences were designed for excision of the region between the first introns of DNAJB1 and PRKACA on human Chromosome 19 using CRISPOR tool (gRNA1 (DNAJB1) 5’-CAGGAGCCGACCCCGTTCGT-3’, gRNA2 (PRKACA): 5’-GTAGACGCGGTTGCGCTAAG-3’).^45^ HepG2 cells were transfected with 10 mg of dual-guide RNAs-CRISPR-CAS9-EGFP vector and Lipofectamine 2000 transfection reagent (Invitrogen, Carlsbad, CA) and incubated at 37 °C and 5% CO2 for 2 days. Single GFP positive cells were sorted at 4°C using a BD FACSAria III (BD Biosciences, Franklin Lakes, NJ) and deposited in 1% gelatin-coated 96 well plates. The excision of the DNAJB1-PRKACA region was confirmed by PCR. The PCR was performed by using a forward primer (DNAJB1-intron 1-F, 5’-AGCTTCTAGCATGTCTGGGG-3’), and a reverse primer (PRKACA-intron 1-R, 5’-CTGGGAAGGCTCATGAGACCT-3’) for DNAJB1-PRKACA fusion site on the genome. Genomic DNA was extracted from clones using Wizard Genomic DNA purification kit (Promega, Madison, WI) and PCR was conducted at a temperature of 98 °C for 5 minutes (1 cycle), 98 °C for 30 seconds, 58 °C for 45 seconds and 72 °C for 1 minute (32 cycles) and 72 °C for 10 minutes (1 cycle) using GoTaq G2 DNA polymerase (Promega). Additionally, DNAJB1-PRKACA fusion mRNA expression was confirmed by reverse-transcription (RT)-PCR. Total RNA was extracted from cells using the Qiagen RNeasy kit (Qiagen, Valencia CA). Complementary DNA was synthesized using High-Capacity cDNA Reverse Transcription Kit (Applied Biosystems, Foster city, CA) as per the manufacturer’s instruction and PCR was carried out at a temperature of 95 °C for 5 minutes (1 cycle), 95 °C for 30 seconds, 56 °C for 30 seconds and 72 °C for 30 seconds (30 cycles) and 72 °C for 5 minutes using GoTaq DNA polymerase (Promega) and a forward primer (hChimera exon 1-F, 5’-GTTCAAGGAGATCGCTGAGG-3), a reverse primer (hChimera exon 3, 5’-TTCCCGGTCTCCTTGTGTTT-3’). The amplified PCR products were isolated and subcloned into pGEM-T easy vector (Promega) and sequenced using Bigdye terminator v3.1 (Applied Biosystems). The Sequence was analyzed by SnapGene software (San Diego, CA).

### Cell Viability Assay

To determine the changes in cell proliferation in HepG2 cells and HepG2^DNAJB1-PRKACA^ clones, we used CellTiter Glo 2 assay systems (Promega). Approximately 2.0 × 10^3^ cells/well were seeded in a 96-well plate and incubated in RPMI1640 culture medium over 4 days at 37 °C and 5% CO_2_. After addition of the CellTiter Glo 2 reagent to cell culture medium in 96 well plate and incubation for 10 minutes, luminescence was measured by CLARIOstar Plus microplate reader (BMG Labtech, Ortenberg, Germany). Additionally, the growth of HepG2 cells and HepG2^DNAJB1-PRKACA^ clones were measured using hemocytometer cell counting. Cells (5.0 ×10^4^ cells/well) were seeded in a 6-well plate and incubated in RPMI1640 culture medium for 4 days at 37 °C and 5% CO_2_. Cells were trypsinized, stained with 0.4% Trypan Blue (Invitrogen) and counted using hemocytometer (Day 0, 2 and 4). For drug-response analysis, HepG2 cells and HepG2^DNAJB1-PRKACA^ clones (H12, H33) (2.0 × 10^3^ cells/well) were seeded in a 96-well plate and incubated with RPMI1640 cell culture medium for 24 hours. After incubation, cells were treated with SY-5609 (1 pM-10 μM) (MedChemExpress, Monmouth Junction, NJ), YKL-5-124 (100 pM-10 μM) (MedChemExpress) or vehicle (DMSO) for 48, 72 hours or 120 hours. Cell viability was measured using CellTiter Glo 2 reagent. Percent of survived cells and growth inhibition were calculated relative to vehicle control (as 100% of survival). The dose-response curves and IC_50_ values were analyzed using GraphPad Prism software (San Diego, CA). Differences between conditions were tested with a t-test and a *p* < 0.05 was taken as statistically significant.

### Caspase 3/7 and 8 assay

To measure the apoptosis activity in HepG2 cells and HepG2^DNAJB1-PRKACA^ clones, we used Caspase-Glo 3/7 and 8 assay systems (Promega). Approximately 2.0 × 10^3^ cells/well were seeded in a 96-well plate and (1) incubated in RPMI1640 culture medium over 4 days at 37 °C and 5% CO_2_ or (2) incubated in RPMI1640 culture medium containing SY-5609 (0.1-10 μM) (MedChemExpress) or DMSO for 24 hours. After addition of the Caspase-Glo 3/7 or 8 reagent to cell culture medium in 96 well plate and incubation for 30 minutes, luminescence was measured by CLARIOstar Plus microplate reader (BMG Labtech).

### Cell Cycle Analysis

To synchronize cell cycle in HepG2 cells and HepG2^DNAJB1-PRKACA^ clones, cells were treated with serum starvation medium (RPMI1640 medium (Life Technologies) supplemented with 1% penicillin– streptomycin-glutamine (Life Technologies)) at 37 °C and 5% CO_2_ for 16 hours. Then serum starvation medium was (1) replaced to RPMI1640 culture medium and cells were collected over 24 hours (every 6 hours) or (2) replaced to RPMI1640 culture medium containing SY-5609 (1 μM) (MedChemExpress) or DMSO for 24 hours and cells were collected. The collected cells were fixed with 70% ethanol and stained with 50 μg/mL propidium iodide (BioLegend, San Diego, CA) and 100 μg/mL RNAse A (Thermo Fisher). Samples were sorted with an Attune NxT flow cytometer (Thermo Fisher) and the DNA content was analyzed for the percent of cells in G0/G1, S, and G2/M phase with ModFit LT 6.0 software at the University of Wisconsin Carbone Cancer Center Flow Cytometry Laboratory. The average coefficient of variation (CV) of each sample was < 6%. Differences between conditions were tested with a t-test and a *p* < 0.05 was taken as statistically significant.

### RNA Isolation, Sequencing, Differential Gene Expression Analysis

We performed RNA isolation and RNA sequencing as previously described.^46^ Briefly, to evaluate for transcriptomic differences between HepG2 and HepG2^DNAJB1-PRKACA^ clones (H33 and H12 cells), bulk RNA-seq was performed on 6 biological replicates collected from each cell line. RNA Isolation was carried out using the RNeasy protocol (Giagen, Hilden, Germany) according to the manufacturer’s recommendations. Quality was tested for an RNA integrity number (RIN) > 7.5 on the Agilent 2100 bioanalyzer (Agilent Technologies, Santa Clara, CA). A total of 300 ng of mRNA was enriched with poly-A selection and sequencing on the Illumina HiSeq2500 platform by the University of Wisconsin Biotechnology Sequencing Core. FASTq files were processed with Skewer and genes were filtered to remove those with low expression.^47^ Samples were normalized by the method of trimmed mean of M-values.^48^ Contrasts were drawn with the edgeR package, with differential expression taken when the FDR *q* < 0.05.^49^ Pathway testing was performed with the KEGG database (Kyoto Encyclopedia of Genes and Genomes) using previously described methods.^50^ The top 500 significant genes were inputted, ordered by *q* value, and the top significant pathways ordered by *q*-value (an adjusted multiple testing p-value found using the false discovery rate approach of Benjamini and Hochberg)^51^ were plotted for visualization. Pathway dot size is indicative of the number of genes in each pathway. All RNA seq data is available at Gene Expression Omnibus (GEO).

### Quantitative reverse transcription PCR (RT-qPCR)

HepG2 and HepG2^DNAJB1-PRKACA^ clones were (1) incubated with RPMI1640 cell culture medium for 24 hours, or (2) treated with SY-5609 (0.1-10 μM) (MedChemExpress), YKL-5-124 (0.1-10 μM) (MedChemExpress) or vehicle (DMSO) for 24 hours. Total RNA was isolated from cells using the Qiagen RNeasy kit (Qiagen, Valencia CA). The isolated RNAs (20 ng) were reverse-transcribed and quantitative PCR (qPCR) were perform using GoTaq Probe 1-Step RT-PCR system (Promega) in a QuantStudio 7 Flex Real-Time PCR System (Applied Biosystems). The reverse-transcription was conducted at a temperature of 45 °C for 15 minutes, and 95 °C for 2 minutes during one cycle. The denaturation, annealing and extension were performed at 95 °C for 15 seconds and 60 °C for 1 minutes (45 cycles). The mRNA levels were measured with Taqman Gene Expression System (FAM-MGB) (Applied Biosystems) (Assay ID: DNAJB1: Hs00356730_g1, DNAJB1-PRKACA fusion: Hs05061318_ft, SLC16A14: Hs00541300_m1, LINC00473: Hs00293257_m1, ETV1: Hs00951951_m1, GATA2: Hs00231119_m1, GATA3: Hs00231122_m1, RUNX1: Hs01021970_m1, SCG2: Hs01920882_s1, CDK7: Hs00361486_m1, GAPDH: Hs02786624_g1, β-Actin: Hs01060665_g1). To detect PRKACA mRNA expression, the qPCR primers (Primer 1: 5’-AGCGGGACTTTCCCATTT-3’, Primer 2: 5’-AAGAAGGGCAGCGAGCA-3’) and FAM-labeled probe (5’-FAM-TGGCTTTGGCTAAGAATTCTTTCACGC-3’) were designed using PrimerQuest Tool (IDT). The qPCR primers and probe spanned the boundary of exon 1-2 junction since DNAJB1-PRKACA fusion gene harbors exon 2-10 of PRKACA gene. GAPDH and β-Actin mRNA levels were used as internal control and the data were analyzed using the ^ΔΔ^Ct method.

### Immunoblotting

HepG2 cells and HepG2^DNAJB1-PRKACA^ clones were lysed in CelLytic M lysis reagent (Sigma-Aldrich) with Halt protease and phosphatase inhibitor cocktail (Thermo Fisher). Human liver homogenate was prepared from human FLC or matched normal liver tissue (adjacent to the FLC) using RIPA Lysis buffer (Thermo Fisher) with Halt protease and phosphatase inhibitor cocktail (Thermo Fisher). For immunoblotting, 20 μg of whole cell extract or 40-80 μg of liver homogenate was loaded on Mini-Protean TGX protein gel (4-20% or 7.5%) (Bio-Rad, Hercules, CA) and transferred to Immobilon-P membranes by electrophoresis (MilliporeSigma, Burlington, MA). Membranes were incubated in 10% w/v BSA blocking buffer (Thermo Fisher) at room temperature for 1 hour and hybridized with primary antibody (PKA(C): 610981 (BD Biosciences), CDK2: #2546 (Cell Signaling Technology), phospho-CDK2 (Thr160): #2561 (Cell Signaling Technology), CDK7: #2916 (Cell Signaling Technology), phospho-CDK7 (T170): ab155976 (abcam, Cambrige, MA), RNA polymerase II RPB-1: RPB-1 NTD; #14958, Phospho-CTD (Ser2); #13499, Phospho-CTD (Ser5); #13523, Phospho-CTD (Ser7); #13780 (Cell Signaling Technology), SCG2; PA5-115018 (Thermo Fisher), PARP; #9542 (Cell Signaling Technology), Cleaved PARP (Asp214); #5625 (Cell Signaling Technology), β-Actin: #4967 (Cell Signaling Technology), MAB8929 (R&D Systems, Minneapolis, MN) at 4°C overnight. After hybridization with Alkaline Phosphatase (AP)**-**conjugated secondary antibody (anti-mouse: #7056 (Cell Signaling Technology), anti-rabbit: 111-055-144 (Jackson ImmunoResearch, West Grove, PA)), proteins were detected using NBT/BCIP solution (Thermo Fisher).

### siRNA transfection

HepG2 cells and HepG2^DNAJB1-PRKACA^ clones were transfected in 96-well plate with 0.25-100 nM Silencer small interfering RNA (siRNA) (Negative Control: #1, CDK7: s2830) (Life Technologies) using Lipofectamine RNAiMAX transfection reagent (Life Technologies). Four days after the transfection, cell viability was measured using CellTiter-Glo 2 reagent (Promega). After addition of the reagent to cell culture medium, luminescence was measured by CLARIOstar Plus microplate reader (BMG Labtech).

### Primary Human Hepatocyte

Primary human hepatocytes (PHHs) were prepared from fresh human liver samples. Liver samples were perfused using perfusion solution I and liver tissues were dissociated by Digestion-Solution containing collagenase P.^52^ PHHs were isolated and purified from hepatocyte rich cell suspension using 25% Percoll solution (Sigma-Aldrich) for density gradient centrifugation. Approximately 2.0 × 10^4^ purified PHH cells/well were seeded in rat tail collagen (collagen type I) (Life Technologies) coated 96-well plate and incubated in hepatocyte Incubation medium at 37 °C and 5% CO_2_.^52^ Twenty-four hours after incubation, PHHs were treated with SY-5609 (100 pM - 10 μM) (MedChemExpress) or vehicle (DMSO) for 2-5 days. Cell viability was detected using CellTiter Glo 2 reagent (Promega). After addition of the reagent to cell culture medium, luminescence was measured by CLARIOstar Plus microplate reader (BMG Labtech).

### FLC Human Cell Line

FLC-H cells were generated from a patient-derived xenograft model (Oikawa et al., 2015) and grown in RPMI1640 media containing 300 mg/L l-glutamine (Thermo Fisher Scientific) supplemented with 10% fetal bovine serum (Thermo Fisher Scientific), 1% penicillin-streptomycin (Thermo Fisher Scientific) and 2.5 μg/mL human hepatic growth factor (Thermo Fisher Scientific). Cells were cultured in a humid chamber at 37°C and 5% CO_2_. FLC1025 and FLCmet3 cells were generated directly from a human FLC tumors and maintained in DMEM/F12 supplemented with 10% FBS (Thermo Fisher Scientific), 0.04 µg/mL dexamethasone (Thermo Fisher Scientific), 0.1% gentamicin (Thermo Fisher Scientific), 1 µg/mL recombinant human insulin (Thermo Fisher Scientific), 0.55 µg/mL human transferrin (Thermo Fisher Scientific), and 0.5 ng/mL sodium selenite (Thermo Fisher Scientific). The media was supplemented with 10% DMEM from cultured human embryonic kidney cells harboring a human RSPO1 transgene and 20 μM Y-27632 ROCK inhibitor. FLC cells were plated at a density of 0.1 x 10^6^ cells/well in 12-well plates. After overnight incubation, cells were treated with vehicle control or SY-5609. Six days post-treatment cells were harvested for counting by hemocytometer. Cell viability was determined using trypan blue staining. Cell counting was performed in three biological replicates per trial and two trials were performed for each condition.

### Tissue slice preparation

FLC tumor slices were prepared as described previously (PMID: 31741771, PMID: 32420994). Briefly, dissected tumor tissues were cut into 400-µm organotypic tumor slices using the Leica VT1200S vibratome microtome (Nusslock) with HBSS as the cutting medium. The slices were then cut into 400-µm cuboids using a McIlwain tissue chopper (Ted Pella) as described previously (PMID: 33174580). Cuboids were immediately placed into 96-well ultralow-attachment plates (Corning) and incubated with Williams’ medium containing 12 mM nicotinamide, 150 nM ascorbic acid, 2.25 mg/mL sodium bicarbonate, 20 mM HEPES, 50 mg/mL additional glucose, 1 mM sodium pyruvate, 2 mM L-glutamine, 1% (v/v) ITS, 20 ng/mL EGF, 40 IU/mL penicillin, and 40 μg/mL streptomycin containing RealTime Glo reagent (Promega) according to the manufacturer’s instructions. After 24 h, the baseline cell viability of cuboids was measured by RealTime Glo bioluminescence using the Synergy H4 instrument (Biotek). Cuboids were exposed to DMSO (control), SY-5609 (0.1 - 1 μM) or Staurosporine (0.5 μM; positive control), and overall tumor tissue viability was measured daily, up to 7 d after treatment.

### Combination Therapy

To investigate the synergistic effect between SY-5609 and A-1331852 (MedChemExpress), HepG2^DNAJB1-PRKACA^ (H33) cells (2.0 × 10^3^ cells/well) were seeded in an 8x8 matrix in a 96-well plate and incubated with RPMI1640 cell culture medium for 24 hours at 37 °C and 5% CO_2_. Cells were treated with SY-5609 (0.1-100 nM) (MedChemExpress) and A-1331852 (0.1-100 nM) (MedChemExpress) or vehicle (DMSO) for 72 hours. Cell viability was measured using CellTiter Glo 2 reagent. Percent of survived cells and growth inhibition were calculated relative to vehicle control (as 100% of survival). Synergy scores and the dose-response curves were analyzed using SynergyFinder tool. Drug synergy was evaluated using the highest single-agent (HSA) model and Loewe model.^53^ The combination effect of SY-5609 and A-1331852 was examined in PHH and HepG2^DNAJB1-PRKACA^ (H33) cells. HepG2^DNAJB1-PRKACA^ (H33) cells (2.0 × 10^3^ cells/well) and PHH (2.0 × 10^4^ cells/well) were seeded in rat tail collagen (collagen type I) coated 96-well plate and incubated in hepatocyte Incubation medium or RPMI1640 cell culture medium at 37 °C and 5% CO_2_ for 24 hours. Cells were treated with SY-5609 (1 μM) (MedChemExpress) and A-1331852 (10 nM) (MedChemExpress) or vehicle (DMSO) for 24 or 48 hours. Cell viability was measured using CellTiter Glo 2 reagent and CLARIOstar Plus microplate reader (BMG Labtech).

### Statistical Analysis

All analyses were performed with GraphPad Prism software (San Diego, CA) unless otherwise indicated. Samples were not excluded from the analysis unless explicitly stated. Standard comparisons of the mean were performed with a t-test with standard deviation unless explicitly stated.

## Notes

### Competing Interest Statement

The authors have declared no competing interest.

### Summary of Updates

I have revised the figures, added new data (including new figures) and revised the manuscript.

