## Supplemental Data 1 for "CDK7 is a Novel Therapeutic Vulnerability in Fibrolamellar Carcinoma"

| Name | Symbol | H12-1 | H12-2 | H12-3 | H12-4 | H12-5 | H12-6 | H33-1 | H33-2 | H33-3 | H33-4 | H33-5 | H33-6 | HepG2-1 | HepG2-2 | HepG2-3 | HepG2-4 | HepG2-5 | HepG2-6 |
|---|---|---|---|---|---|---|---|---|---|---|---|---|---|---|---|---|---|---|---|
| ABL proto-oncogene 1, non-receptor tyrosine kinase | ABL1 | 2381 | 3277 | 2670 | 1927 | 2698 | 2244 | 2334 | 2908 | 2644 | 1912 | 2099 | 2219 | 2297 | 1672 | 1838 | 1412 | 1737 | 1829 |
| anaphase promoting complex subunit 1 | ANAPC1 | 4728 | 5255 | 4272 | 2779 | 4604 | 3697 | 3172 | 3841 | 4224 | 3155 | 3464 | 3185 | 5842 | 4611 | 4791 | 4016 | 4798 | 5061 |
| anaphase promoting complex subunit 10 | ANAPC10 | 252 | 252 | 252 | 143 | 252 | 206 | 182 | 248 | 245 | 170 | 190 | 213 | 319 | 230 | 235 | 210 | 223 | 264 |
| anaphase promoting complex subunit 11 | ANAPC11 | 494 | 802 | 653 | 540 | 681 | 591 | 524 | 900 | 637 | 358 | 485 | 627 | 690 | 531 | 637 | 403 | 608 | 641 |
| anaphase promoting complex subunit 2 | ANAPC2 | 602 | 972 | 730 | 600 | 809 | 658 | 695 | 999 | 749 | 481 | 568 | 734 | 447 | 380 | 334 | 263 | 364 | 381 |
| anaphase promoting complex subunit 4 | ANAPC4 | 823 | 876 | 698 | 525 | 777 | 642 | 556 | 684 | 744 | 580 | 623 | 567 | 878 | 637 | 671 | 577 | 634 | 722 |
| anaphase promoting complex subunit 5 | ANAPC5 | 6083 | 7635 | 6286 | 4592 | 6941 | 5709 | 5406 | 6557 | 6823 | 5057 | 5654 | 5423 | 4627 | 3477 | 4094 | 3213 | 3951 | 4025 |
| anaphase promoting complex subunit 7 | ANAPC7 | 3298 | 4017 | 3309 | 2282 | 3747 | 2900 | 2635 | 3205 | 3270 | 2439 | 2730 | 2572 | 1937 | 1391 | 1629 | 1258 | 1506 | 1581 |
| ATM serine/threonine kinase | ATM | 2367 | 2844 | 2369 | 1454 | 2303 | 1717 | 1753 | 1895 | 1950 | 1651 | 1618 | 1488 | 2279 | 1773 | 1795 | 1643 | 1942 | 2105 |
| ATR serine/threonine kinase | ATR | 2051 | 2451 | 2105 | 1389 | 2164 | 1622 | 1682 | 1754 | 1949 | 1519 | 1536 | 1461 | 1146 | 928 | 896 | 771 | 919 | 983 |
| BUB1 mitotic checkpoint serine/threonine kinase | BUB1 | 5441 | 6267 | 5298 | 3536 | 5837 | 4374 | 4278 | 4970 | 5598 | 4296 | 4705 | 4378 | 3534 | 2666 | 3125 | 2345 | 2915 | 2829 |
| BUB1 mitotic checkpoint serine/threonine kinase B | BUB1B | 3118 | 3524 | 2928 | 2074 | 3270 | 2481 | 2455 | 2866 | 2971 | 2352 | 2589 | 2316 | 2799 | 2193 | 2434 | 1916 | 2265 | 2208 |
| BUB3 mitotic checkpoint protein | BUB3 | 6736 | 7731 | 6757 | 4598 | 7416 | 5767 | 5688 | 6735 | 6943 | 5337 | 5695 | 5568 | 4273 | 3246 | 3606 | 2834 | 3477 | 3656 |
| cyclin A1 | CCNA1 | 0 | 0 | 0 | 0 | 0 | 0 | 0 | 0 | 0 | 0 | 0 | 0 | 0 | 0 | 0 | 0 | 0 | 0 |
| cyclin A2 | CCNA2 | 4333 | 4711 | 4173 | 2765 | 4482 | 3389 | 3234 | 3793 | 4233 | 3105 | 3537 | 3269 | 4687 | 3601 | 3955 | 3056 | 3730 | 3692 |
| cyclin B1 | CCNB1 | 16205 | 19258 | 16719 | 11658 | 18183 | 14148 | 13247 | 15841 | 16628 | 11776 | 13603 | 13163 | 5222 | 4349 | 4896 | 3402 | 4231 | 4211 |
| cyclin B2 | CCNB2 | 3950 | 4642 | 3959 | 2775 | 4372 | 3340 | 3258 | 3928 | 4105 | 2938 | 3405 | 3231 | 1628 | 1301 | 1465 | 1132 | 1380 | 1313 |
| cyclin B3 | CCNB3 | 66 | 84 | 72 | 40 | 73 | 60 | 58 | 69 | 80 | 47 | 63 | 67 | 1 | 2 | 5 | 0 | 2 | 1 |
| cyclin D1 | CCND1 | 3789 | 4120 | 3657 | 2402 | 3709 | 3049 | 2475 | 3280 | 3072 | 2022 | 2446 | 2536 | 8158 | 5934 | 6599 | 5179 | 6331 | 6687 |
| cyclin D2 | CCND2 | 2 | 2 | 2 | 0 | 3 | 1 | 2 | 1 | 2 | 0 | 1 | 0 | 21 | 13 | 17 | 13 | 19 | 15 |
| cyclin D3 | CCND3 | 1008 | 1357 | 1122 | 875 | 1258 | 1113 | 1054 | 1408 | 1243 | 908 | 1028 | 1115 | 246 | 198 | 208 | 191 | 209 | 252 |
| cyclin E1 | CCNE1 | 1201 | 1290 | 1173 | 813 | 1262 | 958 | 905 | 1049 | 1187 | 887 | 1020 | 1000 | 1228 | 845 | 970 | 817 | 968 | 1133 |
| cyclin E2 | CCNE2 | 786 | 908 | 818 | 525 | 910 | 656 | 612 | 705 | 767 | 637 | 673 | 668 | 263 | 174 | 163 | 152 | 180 | 206 |
| cyclin H | CCNH | 1225 | 1382 | 1149 | 795 | 1291 | 984 | 941 | 1217 | 1195 | 981 | 1079 | 1002 | 1217 | 923 | 1015 | 785 | 948 | 1006 |
| cell division cycle 14A | CDC14A | 235 | 232 | 188 | 133 | 218 | 162 | 161 | 166 | 205 | 140 | 183 | 151 | 175 | 149 | 174 | 160 | 167 | 161 |
| cell division cycle 14B | CDC14B | 809 | 658 | 568 | 357 | 556 | 405 | 415 | 460 | 516 | 382 | 422 | 430 | 2429 | 1769 | 1940 | 1736 | 1948 | 2071 |
| cell division cycle 16 | CDC16 | 1529 | 1681 | 1548 | 1095 | 1644 | 1273 | 1174 | 1471 | 1509 | 1180 | 1234 | 1128 | 1372 | 1091 | 1089 | 976 | 1149 | 1210 |
| cell division cycle 20 | CDC20 | 9000 | 12014 | 9111 | 7258 | 10214 | 8918 | 8053 | 11102 | 10609 | 7131 | 8293 | 8768 | 2772 | 2289 | 2577 | 1723 | 2203 | 2215 |
| cell division cycle 23 | CDC23 | 2834 | 3267 | 2740 | 1728 | 2892 | 2279 | 2218 | 2527 | 2737 | 2097 | 2410 | 2074 | 2197 | 1626 | 1738 | 1485 | 1761 | 1902 |
| cell division cycle 25A | CDC25A | 1057 | 976 | 942 | 620 | 970 | 786 | 700 | 929 | 905 | 680 | 737 | 744 | 1927 | 1354 | 1629 | 1345 | 1532 | 1696 |
| cell division cycle 25B | CDC25B | 10097 | 13847 | 11220 | 8369 | 11823 | 10069 | 10623 | 13129 | 12921 | 9292 | 10243 | 10192 | 2137 | 1714 | 1801 | 1299 | 1571 | 1561 |
| cell division cycle 25C | CDC25C | 1306 | 1586 | 1274 | 873 | 1361 | 1062 | 973 | 1258 | 1284 | 1030 | 1115 | 1047 | 1005 | 808 | 839 | 643 | 814 | 743 |
| cell division cycle 26 | CDC26 | 637 | 713 | 670 | 414 | 640 | 538 | 452 | 615 | 616 | 484 | 565 | 494 | 633 | 450 | 491 | 435 | 443 | 487 |
| cell division cycle 27 | CDC27 | 6225 | 6862 | 6091 | 3977 | 6343 | 4804 | 4821 | 5601 | 6095 | 4934 | 5138 | 4811 | 4937 | 3690 | 4141 | 3279 | 3959 | 4093 |
| cell division cycle 6 | CDC6 | 3195 | 3340 | 2983 | 2021 | 3381 | 2648 | 2398 | 2756 | 3052 | 2217 | 2641 | 2496 | 4608 | 3241 | 3626 | 3096 | 3651 | 4225 |
| cell division cycle 7 | CDC7 | 933 | 1100 | 950 | 606 | 1007 | 791 | 767 | 883 | 959 | 780 | 811 | 679 | 398 | 293 | 295 | 281 | 331 | 302 |
| cyclin-dependent kinase 1 | CDK1 | 8285 | 9187 | 7890 | 5366 | 8618 | 6394 | 6407 | 7267 | 8540 | 6780 | 7424 | 6520 | 4445 | 3392 | 3893 | 2959 | 3685 | 3822 |
| cyclin-dependent kinase 2 | CDK2 | 2012 | 2407 | 1953 | 1376 | 2163 | 1775 | 1539 | 1900 | 1947 | 1570 | 1614 | 1536 | 1770 | 1371 | 1398 | 1233 | 1348 | 1552 |
| cyclin-dependent kinase 4 | CDK4 | 5150 | 6026 | 4854 | 3599 | 5540 | 4705 | 4097 | 5690 | 5221 | 3916 | 4420 | 4281 | 5105 | 3898 | 4547 | 3450 | 4350 | 4608 |
| cyclin-dependent kinase 6 | CDK6 | 3839 | 3980 | 3409 | 2055 | 3626 | 2740 | 2869 | 3185 | 3425 | 2685 | 2788 | 2606 | 7451 | 5708 | 5975 | 5195 | 6115 | 6339 |
| cyclin-dependent kinase 7 | CDK7 | 1327 | 1451 | 1182 | 802 | 1374 | 1129 | 961 | 1261 | 1268 | 1011 | 1111 | 965 | 1022 | 755 | 852 | 631 | 796 | 934 |
| cyclin-dependent kinase 9 | CDK9 | 1199 | 1843 | 1466 | 1112 | 1544 | 1307 | 1296 | 1732 | 1541 | 1109 | 1116 | 1312 | 690 | 525 | 631 | 452 | 504 | 584 |
| cyclin-dependent kinase 12 | CDK12 | 4810 | 5831 | 4777 | 3160 | 4912 | 4006 | 3924 | 4676 | 4688 | 3684 | 3818 | 3494 | 5517 | 4142 | 4301 | 3645 | 4339 | 4577 |
| cyclin-dependent kinase 13 | CDK13 | 1924 | 2256 | 1954 | 1191 | 2009 | 1564 | 1487 | 1842 | 1884 | 1537 | 1655 | 1533 | 1164 | 861 | 916 | 798 | 846 | 1001 |
| cyclin-dependent kinase 15 | CDK15 | 290 | 325 | 345 | 218 | 359 | 276 | 281 | 402 | 354 | 201 | 313 | 279 | 1 | 0 | 0 | 0 | 0 | 0 |
| cyclin-dependent kinase 17 | CDK17 | 881 | 938 | 820 | 536 | 909 | 661 | 661 | 781 | 857 | 699 | 739 | 626 | 692 | 483 | 577 | 411 | 570 | 0 |
| cyclin-dependent kinase 20 | CDK20 | 123 | 162 | 148 | 104 | 164 | 114 | 138 | 171 | 146 | 107 | 89 | 121 | 1 | 3 | 2 | 0 | 2 | 2186 |
| cyclin-dependent kinase inhibitor 1A (p21, Cip1) | CDKN1A | 3086 | 3906 | 3119 | 2299 | 3117 | 2813 | 3296 | 4275 | 4120 | 3070 | 3169 | 3451 | 2420 | 1679 | 1850 | 1563 | 1719 | 1412 |
| cyclin-dependent kinase inhibitor 1B (p27, Kip1) | CDKN1B | 1304 | 1417 | 1265 | 863 | 1292 | 982 | 928 | 1129 | 1230 | 841 | 961 | 983 | 1713 | 1278 | 1397 | 1026 | 1473 | 0 |
| cyclin-dependent kinase inhibitor 1C (p57, Kip2) | CDKN1C | 18 | 28 | 34 | 20 | 33 | 17 | 25 | 30 | 30 | 26 | 29 | 21 | 0 | 0 | 0 | 0 | 0 | 32 |
| cyclin-dependent kinase inhibitor 2A | CDKN2A | 3 | 0 | 0 | 0 | 0 | 0 | 0 | 0 | 1 | 0 | 1 | 0 | 25 | 21 | 26 | 18 | 24 | 46 |
| cyclin-dependent kinase inhibitor 2B (p15, inhibits CDK4) | CDKN2B | 6 | 1 | 0 | 0 | 0 | 0 | 5 | 0 | 1 | 1 | 2 | 1 | 48 | 30 | 27 | 35 | 37 | 491 |
| cyclin-dependent kinase inhibitor 2C (p18, inhibits CDK4) | CDKN2C | 1037 | 1329 | 1070 | 745 | 1173 | 958 | 1137 | 1488 | 1394 | 1018 | 1134 | 1228 | 580 | 416 | 501 | 373 | 512 | 105 |
| cyclin-dependent kinase inhibitor 2D (p19, inhibits CDK4) | CDKN2D | 146 | 195 | 157 | 108 | 180 | 125 | 151 | 214 | 182 | 138 | 168 | 157 | 104 | 93 | 90 | 83 | 102 | 1540 |
| checkpoint kinase 1 | CHEK1 | 1603 | 1647 | 1564 | 988 | 1684 | 1244 | 1014 | 1281 | 1350 | 1068 | 1200 | 1068 | 1732 | 1221 | 1436 | 1130 | 1394 | 1267 |
| checkpoint kinase 2 | CHEK2 | 1288 | 1457 | 1228 | 823 | 1326 | 1091 | 1101 | 1313 | 1456 | 1163 | 1252 | 1088 | 1445 | 1161 | 1220 | 954 | 1206 | 1187 |
| CREB binding protein | CREBBP | 1368 | 1997 | 1520 | 1023 | 1462 | 1189 | 1228 | 1513 | 1496 | 1069 | 1162 | 1144 | 1442 | 1146 | 1132 | 932 | 1116 | 2298 |
| cullin 1 | CUL1 | 3278 | 3850 | 3263 | 2133 | 3501 | 2726 | 2528 | 3023 | 3299 | 2535 | 2750 | 2567 | 2737 | 1989 | 2251 | 1831 | 2172 | 820 |
| DBF4 zinc finger | DBF4 | 1721 | 1850 | 1639 | 1011 | 1754 | 1304 | 1189 | 1439 | 1496 | 1136 | 1315 | 1170 | 1096 | 822 | 870 | 656 | 800 | 1884 |
| E2F transcription factor 1 | E2F1 | 1053 | 1278 | 1090 | 813 | 1174 | 1006 | 963 | 1295 | 1141 | 787 | 929 | 991 | 2181 | 1621 | 1851 | 1362 | 1718 | 555 |
| E2F transcription factor 2 | E2F2 | 667 | 834 | 646 | 491 | 750 | 569 | 604 | 819 | 769 | 616 | 620 | 651 | 683 | 511 | 547 | 469 | 538 | 1023 |
| E2F transcription factor 3 | E2F3 | 1874 | 2158 | 1920 | 1215 | 1988 | 1595 | 1293 | 1632 | 1751 | 1319 | 1435 | 1308 | 1220 | 915 | 922 | 768 | 949 | 2447 |
| E1A binding protein p300 | EP300 | 2832 | 3600 | 2710 | 1853 | 2805 | 2270 | 2290 | 2615 | 2660 | 2118 | 2126 | 2032 | 2968 | 2252 | 2268 | 2110 | 2325 | 757 |
| extra spindle pole bodies homolog 1 (S. cerevisiae) | ESPL1 | 1570 | 2329 | 1670 | 1249 | 1742 | 1511 | 1503 | 1994 | 1806 | 1322 | 1425 | 1520 | 942 | 731 | 775 | 566 | 757 | 407 |
| fizzy/cell division cycle 20 related 1 (Drosophila) | FZR1 | 616 | 933 | 708 | 616 | 724 | 687 | 690 | 1011 | 757 | 541 | 622 | 752 | 528 | 402 | 448 | 290 | 401 | 434 |
| growth arrest and DNA-damage-inducible, alpha | GADD45A | 478 | 599 | 512 | 321 | 521 | 430 | 352 | 525 | 528 | 395 | 406 | 439 | 662 | 430 | 480 | 413 | 487 | 171 |
| growth arrest and DNA-damage-inducible, beta | GADD45B | 177 | 218 | 180 | 130 | 193 | 175 | 154 | 168 | 194 | 89 | 158 | 147 | 281 | 192 | 221 | 165 | 214 | 0 |
| growth arrest and DNA-damage-inducible, gamma | GADD45G | 1 | 1 | 1 | 0 | 0 | 0 | 1 | 0 | 1 | 2 | 0 | 0 | 0 | 0 | 0 | 0 | 0 | 1590 |
| glycogen synthase kinase 3 beta | GSK3B | 2166 | 2548 | 2139 | 1318 | 2307 | 1761 | 1754 | 1960 | 2121 | 1743 | 1788 | 1653 | 1849 | 1355 | 1470 | 1256 | 1401 | 3916 |
| histone deacetylase 1 | HDAC1 | 10494 | 12393 | 10055 | 7336 | 11280 | 9143 | 8965 | 11269 | 11503 | 8756 | 9338 | 9088 | 4694 | 3517 | 4002 | 3033 | 3827 | 3623 |
| histone deacetylase 2 | HDAC2 | 4875 | 5289 | 4763 | 3226 | 5125 | 4128 | 3793 | 4484 | 4858 | 3726 | 3990 | 3724 | 4254 | 3153 | 3538 | 2859 | 3338 | 488 |
| MAD1 mitotic arrest deficient-like 1 (yeast) | MAD1L1 | 612 | 884 | 651 | 443 | 685 | 615 | 576 | 929 | 758 | 513 | 564 | 676 | 726 | 479 | 544 | 362 | 528 | 2588 |
| MAD2 mitotic arrest deficient-like 1 (yeast) | MAD2L1 | 3653 | 4124 | 3783 | 2480 | 4118 | 3105 | 2794 | 3515 | 3724 | 2817 | 3135 | 2790 | 3117 | 2428 | 2690 | 2035 | 2453 | 917 |
| MAD2 mitotic arrest deficient-like 2 (yeast) | MAD2L2 | 1307 | 1878 | 1368 | 1134 | 1530 | 1375 | 1248 | 1777 | 1636 | 1167 | 1212 | 1461 | 1089 | 842 | 949 | 739 | 907 | 3494 |
| minichromosome maintenance complex component 2 | MCM2 | 5034 | 7276 | 5854 | 4391 | 6099 | 5427 | 4602 | 6544 | 5566 | 3847 | 4519 | 5021 | 4112 | 3048 | 3350 | 2522 | 3340 | 10729 |
| minichromosome maintenance complex component 3 | MCM3 | 6678 | 6969 | 5808 | 4034 | 6380 | 5213 | 4244 | 5270 | 5268 | 3757 | 4384 | 4207 | 12425 | 9250 | 10124 | 8681 | 10415 | 6711 |
| minichromosome maintenance complex component 4 | MCM4 | 8460 | 10037 | 8741 | 5796 | 9186 | 7589 | 6565 | 8072 | 8085 | 5951 | 6828 | 6659 | 7816 | 5562 | 6085 | 5282 | 6343 | 2973 |
| minichromosome maintenance complex component 5 | MCM5 | 1920 | 2713 | 2147 | 1736 | 2307 | 2091 | 1606 | 2385 | 2008 | 1291 | 1596 | 1855 | 3462 | 2517 | 2778 | 2048 | 2853 | 2429 |
| minichromosome maintenance complex component 6 | MCM6 | 6147 | 6991 | 6216 | 4319 | 7021 | 5430 | 4826 | 5882 | 5882 | 4465 | 5016 | 4913 | 2572 | 1937 | 2081 | 1763 | 2210 | 8376 |
| minichromosome maintenance complex component 7 | MCM7 | 13717 | 17306 | 14204 | 10674 | 15903 | 13987 | 10911 | 14657 | 13982 | 9837 | 11583 | 12155 | 9748 | 7417 | 8194 | 6652 | 8131 | 6629 |
| MDM2 proto-oncogene, E3 ubiquitin protein ligase | MDM2 | 2496 | 2202 | 1719 | 1125 | 1819 | 1437 | 1739 | 1921 | 1834 | 1567 | 1536 | 1447 | 8164 | 5987 | 6049 | 5125 | 6077 | 10554 |
| proliferating cell nuclear antigen | PCNA | 6397 | 6548 | 6013 | 3964 | 6435 | 5182 | 4412 | 6210 | 5670 | 4260 | 5064 | 4832 | 12029 | 8728 | 9571 | 7923 | 9605 | 200 |
| protein kinase, membrane associated tyrosine/threonine 1 | PKMYT1 | 162 | 315 | 254 | 196 | 247 | 206 | 244 | 352 | 211 | 161 | 199 | 237 | 281 | 182 | 251 | 146 | 185 | 3456 |
| polo-like kinase 1 | PLK1 | 4177 | 5470 | 4311 | 3330 | 4480 | 3844 | 3751 | 4832 | 4593 | 3156 | 3592 | 4116 | 4552 | 3644 | 4049 | 2754 | 3541 | 8978 |
| protein kinase, DNA-activated, catalytic polypeptide | PRKDC | 16550 | 23240 | 17519 | 10784 | 18060 | 13760 | 13287 | 16844 | 15802 | 13205 | 13106 | 12050 | 9352 | 7077 | 6835 | 7116 | 8307 | 1293 |
| pituitary tumor-transforming 1 | PTTG1 | 5042 | 5976 | 5085 | 3712 | 5780 | 4303 | 4346 | 5215 | 5490 | 3863 | 4507 | 4234 | 1625 | 1332 | 1570 | 1090 | 1359 | 0 |
| pituitary tumor-transforming 2 | PTTG2 | 0 | 0 | 1 | 0 | 0 | 0 | 0 | 0 | 0 | 0 | 0 | 0 | 0 | 0 | 0 | 0 | 0 | 1717 |
| retinoblastoma 1 | RB1 | 3179 | 3640 | 3142 | 2095 | 3138 | 2484 | 2381 | 2753 | 3007 | 2339 | 2587 | 2362 | 2042 | 1439 | 1574 | 1410 | 1660 | 1471 |
| retinoblastoma-like 1 | RBL1 | 1196 | 1276 | 1036 | 740 | 1182 | 903 | 867 | 1002 | 1059 | 902 | 929 | 894 | 1718 | 1233 | 1387 | 1159 | 1389 | 2232 |
| retinoblastoma-like 2 | RBL2 | 4618 | 5266 | 4371 | 2997 | 4606 | 3595 | 3421 | 3965 | 4037 | 3251 | 3386 | 3307 | 2773 | 2013 | 2349 | 1766 | 2269 | 1139 |
| ring-box 1, E3 ubiquitin protein ligase | RBX1 | 1033 | 1119 | 952 | 706 | 1076 | 813 | 759 | 945 | 1038 | 692 | 910 | 866 | 1308 | 980 | 1160 | 874 | 1118 | 293 |
| stratifin | SFN | 135 | 206 | 125 | 125 | 121 | 125 | 161 | 273 | 223 | 148 | 143 | 249 | 410 | 279 | 304 | 210 | 287 | 5297 |
| S-phase kinase-associated protein 1 | SKP1 | 8125 | 8751 | 7974 | 5200 | 8891 | 6820 | 6095 | 7812 | 8019 | 6109 | 6663 | 6346 | 6091 | 4712 | 5176 | 4182 | 4985 | 2670 |
| S-phase kinase-associated protein 2, E3 ubiquitin protein ligase | SKP2 | 2037 | 2185 | 1811 | 1216 | 1981 | 1539 | 1459 | 1748 | 1836 | 1388 | 1531 | 1508 | 3209 | 2385 | 2484 | 2085 | 2597 | 1554 |
| SMAD family member 2 | SMAD2 | 2295 | 2685 | 2293 | 1549 | 2423 | 1986 | 1938 | 2245 | 2470 | 2041 | 2005 | 1865 | 1861 | 1489 | 1468 | 1160 | 1364 | 1983 |
| SMAD family member 3 | SMAD3 | 2886 | 3526 | 2876 | 2063 | 3059 | 2597 | 2064 | 2470 | 2660 | 1900 | 2089 | 1991 | 2500 | 1917 | 1934 | 1626 | 1912 | 1664 |
| SMAD family member 4 | SMAD4 | 1562 | 1692 | 1380 | 947 | 1475 | 1209 | 1145 | 1312 | 1420 | 1117 | 1135 | 1028 | 1942 | 1563 | 1574 | 1254 | 1541 | 4997 |
| structural maintenance of chromosomes 1A | SMC1A | 4836 | 5819 | 4602 | 3040 | 4739 | 3996 | 3745 | 4521 | 4751 | 3743 | 3882 | 3643 | 5900 | 4386 | 4687 | 4065 | 4802 | 2 |
| structural maintenance of chromosomes 1B | SMC1B | 18 | 16 | 11 | 9 | 15 | 15 | 9 | 6 | 9 | 7 | 17 | 12 | 0 | 0 | 2 | 1 | 1 | 3604 |
| transcription factor Dp-1 | TFDP1 | 5137 | 5949 | 4994 | 3642 | 5447 | 4633 | 4015 | 5056 | 5351 | 3893 | 4388 | 4304 | 4325 | 3271 | 3485 | 2782 | 3331 | 735 |
| transforming growth factor, beta 1 | TGFB1 | 801 | 1246 | 914 | 771 | 971 | 871 | 972 | 1383 | 1189 | 855 | 991 | 1143 | 929 | 624 | 682 | 525 | 645 | 1 |
| transforming growth factor, beta 2 | TGFB2 | 4 | 0 | 2 | 5 | 4 | 2 | 2 | 1 | 3 | 0 | 2 | 1 | 0 | 2 | 0 | 1 | 2 | 28 |
| transforming growth factor, beta 3 | TGFB3 | 59 | 75 | 64 | 51 | 63 | 42 | 46 | 69 | 59 | 34 | 64 | 34 | 22 | 16 | 14 | 14 | 21 | 1501 |
| tumor protein p53 | TP53 | 5113 | 6892 | 5744 | 4269 | 5936 | 5103 | 4683 | 5819 | 5577 | 3960 | 4456 | 4595 | 1679 | 1309 | 1493 | 1202 | 1432 | 1695 |
| WEE1 G2 checkpoint kinase | WEE1 | 1525 | 1702 | 1427 | 1042 | 1560 | 1172 | 1091 | 1363 | 1439 | 1133 | 1248 | 1174 | 1993 | 1578 | 1578 | 1369 | 1589 | 10499 |
| tyrosine 3-monooxygenase/tryptophan 5-monooxygenase activation protein, beta | YWHAB | 12568 | 13595 | 11895 | 8095 | 13066 | 10306 | 9316 | 11076 | 11769 | 8700 | 9716 | 9255 | 11765 | 8884 | 9818 | 8070 | 9438 | 9887 |
| tyrosine 3-monooxygenase/tryptophan 5-monooxygenase activation protein, epsilon | YWHAE | 23075 | 27490 | 24091 | 16967 | 26429 | 21085 | 20384 | 23746 | 24791 | 17904 | 20562 | 20026 | 11959 | 8668 | 9685 | 7823 | 9139 | 6025 |
| tyrosine 3-monooxygenase/tryptophan 5-monooxygenase activation protein, gamma | YWHAG | 18007 | 23283 | 19729 | 13498 | 20629 | 16749 | 16755 | 19960 | 20915 | 15968 | 16660 | 16700 | 7257 | 5278 | 5860 | 4571 | 5535 | 2623 |
| tyrosine 3-monooxygenase/tryptophan 5-monooxygenase activation protein, eta | YWHAH | 2847 | 3245 | 2919 | 1892 | 3107 | 2459 | 2231 | 2576 | 2931 | 2141 | 2512 | 2251 | 3033 | 2306 | 2608 | 2003 | 2270 | 10906 |
| tyrosine 3-monooxygenase/tryptophan 5-monooxygenase activation protein, theta | YWHAQ | 24387 | 27344 | 24959 | 16799 | 27773 | 21276 | 19260 | 23298 | 24458 | 18544 | 20837 | 20115 | 12727 | 9599 | 10547 | 8457 | 10033 | 13257 |
| tyrosine 3-monooxygenase/tryptophan 5-monooxygenase activation protein, zeta | YWHAZ | 25048 | 28174 | 25513 | 16894 | 27492 | 21314 | 19629 | 23013 | 24374 | 18734 | 20721 | 19191 | 15703 | 11361 | 12809 | 10487 | 12408 | |
